# Time-course in attractiveness of pheromone lure on the smaller tea tortrix moth: a generalized additive mixed model approach

**DOI:** 10.1101/2020.11.06.370809

**Authors:** Masaaki Sudo, Yasushi Sato, Hiroshi Yorozuya

## Abstract

Long-term pest insect monitoring in agriculture and forestry has advanced population ecology. However, the discontinuation of research materials such as pheromone lure products jeopardizes data collection continuity, which constrains the utilization of the industrial datasets in ecology. Three pheromone lures against the smaller tea tortrix moth *Adoxophyes honmai* Yasuda (Lepidoptera; Tortricidae) were available but one was recently discontinued. Hence, a statistical method is required to convert data among records of moths captured with different lures. We developed several generalized additive mixed models (GAMM) separating temporal fluctuation in the background male density during trapping and attenuation of lure attractiveness due to aging or air exposure after settlement. We collected multisite trap data over four moth generations. The lures in each of these were unsealed at different times before trap settlement. We used cross-validation to select the model with the best generalization performance. The preferred GAMM had nonlinear density fluctuation terms and lure attractiveness decreased exponentially after unsealing. The attenuation rates varied among lures. A light trap dataset near the pheromone traps was a candidate for a male density predictor. Nevertheless, there was only a weak correlation between trap yields, suggesting the difficulty of data conversion between the traps differing in attraction mechanisms.

## 1. Introduction

Long-term field monitoring of population fluctuations is a fundamental information source in population ecology because an aim of the discipline is to clarify the dynamics driving the population of an organism (Reinke, Miller, & Janzen, 2019; Royama, 1996; Yamamura et al., 2006). Large-scale and long-term data have been collected for agriculture, fishery (Jorgensen et al., 2007), hunting (Elton & Nicholson, 1942) and forestry (Klapwijk, Csóka, Hirka, & Björkman, 2013; Osada et al., 2018). Data sources in the industrial sectors have advocated and tested basic concepts in ecology and evolutionary and conservation biology.

However, it has also been difficult to obtain census data with sufficient length and methodological consistency (Bonebrake, Christensen, Boggs, & Ehrlich, 2010). For insect pest species, data recycling has been constrained by temporal changes in chemical pesticides and cultivars (Osakabe, 1985; Yamamura et al., 2006), suspension and/or site change during the census (Yamamura et al., 2006), and data crudeness and ordinal measurements (Osada et al., 2018; Sudo, Yamanaka, & Miyai, 2019). A census dataset may also be inconsistent because of changes in the equipment and materials or the coexistence of multiple materials. To analyze population dynamics using datasets of the pest species across time periods and regions, a statistical framework is needed to derive the background population sizes from the raw trapping data by absorbing differences in research materials.

Since the 1960s, numerous female insect pest sex pheromones have been identified. Synthetic pheromone products have been deployed to control population density via mass trapping and/or mating disruption (Cardé, 1976; Witzgall, Kirsch, & Cork, 2010). Those pheromones have also been utilized to monitor pest insect densities as the attractant of a trap. However, a part of the products were discontinued because of corporate mergers and/or product portfolio management.

The smaller tea tortrix *Adoxophyes honmai* Yasuda (Lepidoptera; Tortricidae) is a common tea pest insect in Japan. It is characterized by four (Honshu Island) or five (Southern Japan) discrete generations (Ishijima, Sato, & Ohtaishi, 2009; Nabeta, Nakai, & Kunimi, 2005; Osakabe, 1986; Sato, Takeda, Onoda, & Takaoka, 2005) per season. Field monitoring of *A. honmai* has been routinely conducted by Japanese agricultural researchers since the end of World War II (Oba, 1979; Osakabe, 1985) because the timing of insecticide application to the larvae is determined from the adult emergence of the previous generation. These legacy data have been used in population ecology analyses (Yamanaka, Nelson, Uchimura, & Bjørnstad, 2012).

The female *A. honmai* sex pheromone was first identified by Tamaki, Noguchi, Yushima, & Hirano (1971) though early *A. honmai* censuses relied solely on fluorescent light traps. It comprises (Z)-9-tetradecenyl acetate (Z9-14Ac) and (Z)-11-tetradecenyl acetate (Z11-14Ac) and causes sexual stimulation in males. (E)-11-tetradecenyl acetate (E11-14Ac) and 10-methyldodecyl acetate (10-Me-12Ac) were later isolated and deemed essential for male moth flight orientation. The mass ratio of the components in the female moths was 63:31:4:2 (Z9-14Ac:Z11-14Ac:E11-14Ac:10-Me-12Ac) (Tamaki et al., 1979).

Three synthetic pheromone products have since been dedicated to monitoring *A. honmai* (Japan Plant Protection Association, 2000; Oba, 1979). Each one has a unique mixing ratio, total component content, and pheromone dispenser substrate. The first was released from Ohtsuka Pharmaceutical Co. Ltd. (hereafter, OT). It has a 23:10:0.1:67 (23%, 10%, 0.1%, and 67%) mixing ratio and its format is 9.1 mg per plastic capsule. The ratio of the fourth component was 30× greater than that occurring in living females. It was used in monitoring programs in several prefectures, but its production was discontinued in 2019. The product from Sumitomo Chemical Co. Ltd. (hereafter, SC; originally developed by the agrochemical division of Takeda Pharmaceutical) has a 15:8:1:9 (45%, 24%, 3%, and 27%) mixing ratio and its format is 10 mg per rubber septum. The third product was commercialized by Shin-Etsu Chemical Co. Ltd. (hereafter, SE). Its unofficial mixing ratio is 65:35:5:200 (21%, 11%, 1.6%, and 66%) and its format is 3.05 mg per rubber septum (Yoshioka & Sakaida, 2008).

Here, we sought an appropriate statistical model structure to separate the effects of temporal fluctuations in population size and the pheromone lure attractiveness. To this end, we used a trap dataset over four adult *A. honmai* generations, three lure types, and daily light trap capture simultaneously. The model structure was decomposed to (1) the state of the system or temporal male population size fluctuation including daily changes in flight activity and (2) the observation process or lure attractiveness and trap capture. We could also assume a parametric formulation considering male population dynamics which ecologists recognize as a state-space model (Fukaya, 2016; Kéry & Schaub, 2011; Yamamura, 2016). For the sake of simplicity, however, we adopted a nonlinear regression.

Along with lure product (OT, SC, and SE) chemical composition, attenuation of the volatile concentrations in each lure with air exposure time affects lure attractiveness (Butler & McDonough 1979; Tamaki & Noguchi, 1983). This change is temporal as are population density and male activity. We must design the model so that it can divide these effects. We compared the yields from traps with different lure ages. This configuration was set up by unsealing some of the lures before the start of the session (Tamaki et al., 1980). Besides, as a way to trap insects independently of lure degradation, we also approximated the moth population size using the number of males captured by a light trap near the pheromone lures.

## 2. Methods

### Experimental setup

The field survey was conducted in the experimental fields of Kanaya Tea Research Station, Shizuoka, Japan (34°48’26.6” N, 138°07’57.5” E, 202 m a.s.l.) (Figure 1a). The 0^th^, 1^st^, 2^nd^, and 3^rd^ *A. honmai* generations usually occur annually in Kanaya (Ishijima, Sato, & Ohtaishi, 2009). We captured the generations 1–3 of the 2017 season and the 0^th^ (overwintering) generation of 2018 using traps equipped with pheromone lures.

**Fig 1.**
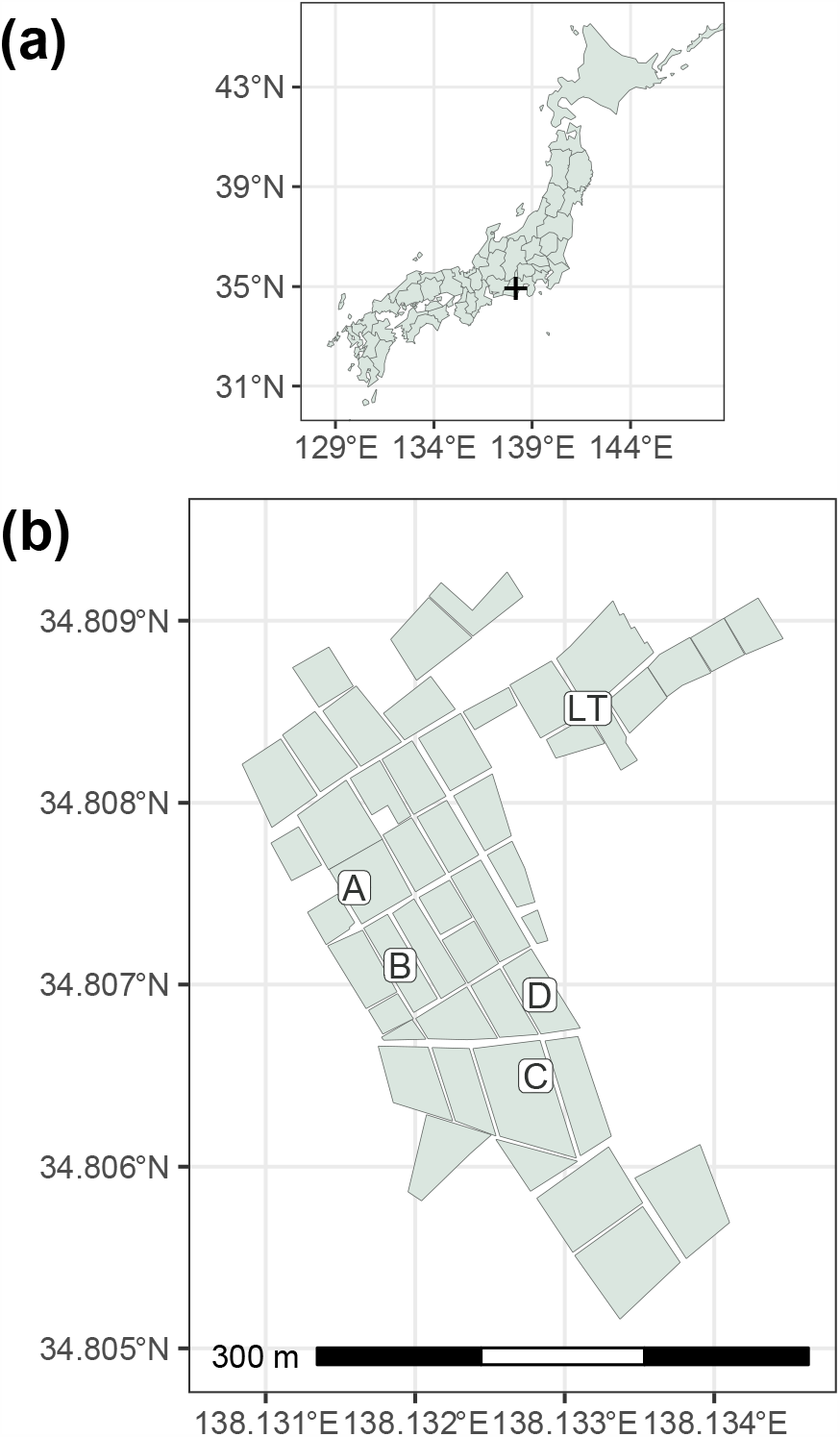
Research field map. Sites “A”–”D” in panel (b) show fields in Kanaya Tea Research Station wherein pheromone traps were settled. Each polygon signifies a tea field. “LT” is the permanent light trap site. Detailed spatial treatment arrangements in tea fields are shown in Supplementary Figure S1.

Each pheromone trap consisted of an adhesive (SE trap; Sankei Chemical Co. Ltd., Kagoshima, Japan) upon which the plastic capsule or the rubber septum of the pheromone lure was placed. The pheromone traps were set in the middle of the tea field hedges at the height of the plucking surface. To calibrate the lure attractiveness based on the number of moths captured per light trap per day, light trap data (Figure 1b: site “LT”) were added to some of the following analyses. Tea-pest monitoring at the light trap site has been conducted since 1947 (Osakabe, 1986). The light trap consisted of a water pan (H 10 cm × W 100 cm × L 100 cm) to capture the insects and a suspended blue fluorescent lamp (20 W; FL20S-BL-K; Panasonic Corp., Osaka, Japan). The light was switched on every night between 19:00 and 7:00 and captured insects including *A. honmai* were counted during the daytime.

The trapping session for each generation was planned to start immediately after the appearance of adults in the field, which was confirmed by the light trap. The pheromone lures were set in the traps on the first day of the session (hereafter, Session Day = 0). Adult male *A. honmai* were counted in the trap every morning from the next day (first day of counting; Session Day = 1). The sessions occurred from June 26, 2017 to July 10, 2017 for the first generation and counting started on June 27, 2017. The second-generation session ran from July 31, 2017 to August 21, 2017. The third-generation session was from September 11, 2017 to October 2, 2017. The zeroth-generation session was from April 28, 2018 to May 12, 2018 (Table 1). The weather at the research station was permanently monitored and meteorological data are available at https://www.naro.affrc.go.jp/org/vegetea/mms-kanaya/Menu.html.

**Table 1.** Weather conditions during the trap sessions were obtained from meteorological data generated by the Kanaya Tea Research Station. There were no observations between July 31, 2017 and August 6, 2018. Hence, data for the 2^nd^ session covers August 7, 2017 to August 21, 2017.

| Generation | 1st | 2nd | 3rd | 0th |
| --- | --- | --- | --- | --- |
| Trap start (SessionDay = 0) | 2017-06-26 | 2017-07-31 | 2017-09-11 | 2018-04-28 |
| Trap end | 2017-07-10 | 2017-08-21 | 2017-10-02 | 2018-05-12 |
| Mean temperature (°C) | 23.7 | 25.3 | 21.3 | 16.1 |
| Min temperature (°C) | 18.3 | 21.9 | 14.4 | 7.6 |
| Max temperature (°C) | 33.3 | 35.1 | 29.9 | 25.5 |
| Precipitation (total) (mm) | 39.5 | 78.0 | 123.5 | 270.5 |
| Precipitation (daily max) (mm) | 26.5 | 37.0 | 83.0 | 82.5 |
| Mean wind velocity (m/s) | 2.9 | 2.6 | 3.0 | 3.4 |
| Max wind velocity (m/s) | 8.2 | 6.6 | 8.6 | 10.3 |
| Mean daylight hours (hour) | 4.6 | 3.3 | 3.9 | 7.3 |
| Mean daily irradiation (MJ/m<sup>2</sup>) | 13.1 | 10.4 | 10.6 | 14.8 |

To separate the temporal changes in the post-settlement lure attractiveness and the population density and activity, the pheromone lures were unsealed at five different times per moth generation. Lures were opened immediately before (Session Day = 0; hereafter, Minus0w) in addition to 7, 14, 21, and 28 days before trap settlement (Minus1w, Minus2w, Minus3w, and Minus4w). The capture session of each generation continued for two (0^th^ and 1^st^) or three weeks (2^nd^ and 3^rd^). Hence, the capture data covered 1–49 (or 1–42 for 0^th^ and 1^st^ generations) days after lure opening.

The experiment was conducted in four fields (sites A–D; Figures 1b and S1) over each of the four moth generations. Each replicate was a combination of pheromone traps distributed within a single tea field and comprised three lure types × five open timings (Supplementary Figure S1). The minimum distance between adjacent pheromone traps was 7.65 m; the attractiveness range for *A. honmai* is ∼5 m (Kawasaki et al., 1979). Hence, interference might have occurred. Nevertheless, an earlier study compared spatiotemporal dynamics in tea tortrix moth lure attractiveness and reported a consensus of 5 m distance between traps (Kawasaki & Tamaki, 1980).

### Model

The model components include the daily male density at each trap site and the attractiveness of the pheromone lure.

#### Male density

Let *d*_*g,i,τ*_ be the density of active males of the *g*^th^ insect generation at the *i*^th^ trap site on the *τ*^th^ trap night. In the scope of the present study, daily male density is defined as the mixture reflecting population dynamics such as recruitment or survival and daily flight activity. It will be affected by weather conditions and male mating drive. The variable *d*_*g,i,τ*_ was expressed as the natural logarithm because the internal population state was here treated as a black box. However, the aforementioned factors synergistically influenced the observed number of moths.

The mean population density of the site *d*_*g,i*_ was determined first. Let *d*_*g*_ be the regional mean (log-) density of *A. honmai* adult males in the *g*^th^ generation. Considering stochastic changes in male density in each tea field, the mean density at the *i*^th^ site during the *g*^th^ generation is:

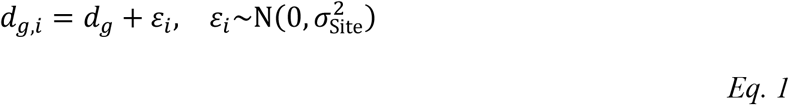

where 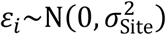 is the random effect of each trap site. It follows a normal distribution with mean = 0 and 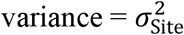. When the trap session of the *g*^th^ generation continues over *T*_*g*_ days, the site-specific male density of the *τ*^th^ day, *d*_*g,i,τ*_ (*τ* = 1,2,3, …, *T*_*g*_), is expressed as:

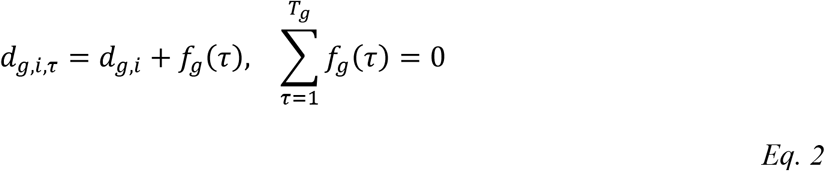

The shape of the generation-specific smoothing function *f*_*g*_(*τ*), signifies the temporal fluctuation in male density during the season. The function is centered about the mean (log-) generation density.

When the male density is approximated by the number captured in the light trap on the same day, the predictor of Eq. 2 should be defined as follows:

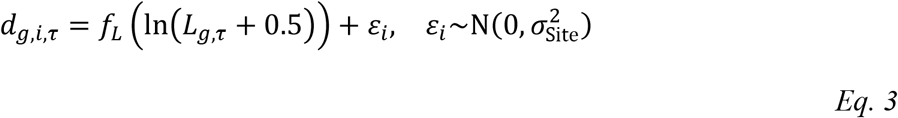

The variable *L*_*g,τ*_ is the measured number of *A. honmai* males captured in the light trap on the same day. As this number may be zero, however, 0.5 is added to it and it is then converted to a logarithm to meet the homoscedasticity assumption (Yamamura, 1999). The shape of the smoothing function *f*_*L*_ indicates the relationship between the numbers caught in the light traps and the pheromone traps. The shape of *f*_*L*_ may be linear or nonlinear and may differ with generation.

#### Lure attractiveness

Next, the lure attractiveness is modeled. As some of the pheromone lures were unsealed before the start of the session, each lure had an age of (pre-session opening period) + (days in the session). Let the lures with the *k*^th^ treatment level (Minus0w, Minus1w, …) age for *m*_*k*_ d at settlement in the fields (Session Day = 0). The lures aged *s*_*k,τ*_ = *τ* + *m*_*k*_ days upon counting Session Day = *τ*. For instance, the first treatment level in the experiment was Minus0w; hence, *m*_*k*=1_ = 0 holds. When the males were counted in the trap for the first time, *s*_*k*=1,*τ* =1_ = 1 holds as *τ* = 1 was defined as the next morning of the trap settlement. If the treatment was Minus1w (*k* = 2), then *m*_*k*=2_ = 7 and *s*_*k*=2,*τ* =1_ = 8.

Let us define the predictor of the number of males captured in each trap when we used the pheromone lure of the *j*^th^ company. Let *μ*_*g,i,j,k,τ*_ be the expected number of unique individuals entering the trap sampling range (site *i*, lure company *j*, and treatment *k*) during the period from *τ* − 1 to *τ* in the trap session of the *g*^th^ generation. Note that *μ*_*g,i,j,k,τ*_ is on a linear scale. When *d*_*g,i,τ*_ gives the local male density, the captured number of males in the morning of Session Day = *τ* is denoted by *N*_*g,i,j,k,τ*_ and is expressed as

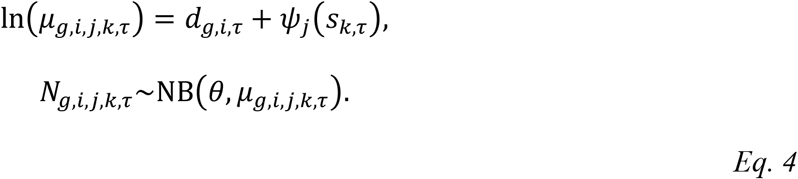

NB(*θ, μ*_*g,i,j,k,τ*_) is the negative binomial distribution of E(*N*_*g,i,j,k,τ*_) = *μ*_*g,i,j,k,τ*_ and 1⁄*θ* is the dispersion parameter such that var(*N*_*g,i,j,k,τ*_) = *μ*_*g,i,j,k,τ*_ + (*μ*_*g,i,j,k,τ*_)^2^ /*θ* (Agresti, 2002).

The effect of the number of days since lure opening (*s*_*k,τ*_) on lure attractiveness is expressed using the continuous function *ψ*_*j*_. Whether the effect of the exposure period is linear or nonlinear, the intercept of *ψ*_*j*_ in Eq. 4 must be fixed for model identifiability. When it is assumed that *ψ*_*j*_(*s*_*k,τ*_) is a linear function, it is conveniently fixed as *ψ*_*j*=1_(0) = 0, which corresponds to the attractiveness of the reference class lure (OT in this study) immediately after unsealing. If it assumed that *ψ*_*j*_(*s*_*k,τ*_) is nonlinear, then it should be centered for the entire session counting (*τ* = 1,2,3,…, *T*_*g*_) as it was for *f*_*g*_(*τ*).

### Parameter estimation

The aforementioned model structure is a type of generalized additive mixed model. To estimate the parameters with the *A. honmai* data, R version 3.6.1 (R Core Team, 2019) and the R package “mgcv” version 1.8-33 (Wood, 2020) were used. To determine whether the shapes of smoothing terms such as *f*_*g*_(*τ*) and *ψ*_*j*_(*s*_*k,τ*_) should be approximated as linear or nonlinear functions, 32 candidate models were constructed and their generalization performances were compared by cross-validation.

#### Models using pheromone traps with different lure ages

A family of candidate models was constructed based on Eq. 2 and Eq. 4. Half of the 32 models (Table 2, models 1–16) were characterized according to whether or not 1) the population fluctuation was linear, 2) the population density intercept was common for all insect generations, 3) the shape of the temporal fluctuation or the slope of the linear variants was common for all insect generations, and 4) the temporal change in lure attractiveness or attenuation after opening was linear.

**Table 2.** Candidate model cross validation. Each model was fitted using a dataset for three of four sites such as A, B, and C. Prediction was then made for the excluded site such as D. This procedure were repeated for all four sites. Goodness-of-fit was the sum of the negative log likelihood (NLL) for the observed number of males at the excluded site. The Models 1 and 2 NLL columns are underlined and signify the best and second-best fits, respectively.

| Model | Formula<br>* | Base density fluctuation of males |  |  |  | Lure<br>attenuation<br>linear? | NLL<br>sum |
| --- | --- | --- | --- | --- | --- | --- | --- |
|  |  | Approximated<br>with light trap? | Linear base<br>density? | Intercept<br>common<br>between<br>generations? | Shape (slope)<br>common<br>between<br>generations? |  |  |
| 1 | I + P | - | - | - | - | - | <u>32669</u> |
| 2 | I + Q | - | - | - | - | TRUE | <u>32555</u> |
| 3 | II + P | - | - | - | TRUE | - | 42709 |
| 4 | II + Q | - | - | - | TRUE | TRUE | 42622 |
| 5 | III + P | - | - | TRUE | - | - | 34057 |
| 6 | III + Q | - | - | TRUE | - | TRUE | 33909 |
| 7 | IV + P | - | - | TRUE | TRUE | - | 45048 |
| 8 | IV + Q | - | - | TRUE | TRUE | TRUE | 44926 |
| 9 | i + P | - | TRUE | - | - | - | 40345 |
| 10 | i + Q | - | TRUE | - | - | TRUE | 40425 |
| 11 | ii + P | - | TRUE | - | TRUE | - | 45773 |
| 12 | ii + Q | - | TRUE | - | TRUE | TRUE | 45758 |
| 13 | iii + P | - | TRUE | TRUE | - | - | 45304 |
| 14 | iii + Q | - | TRUE | TRUE | - | TRUE | 45337 |
| 15 | iv + P | - | TRUE | TRUE | TRUE | - | 48072 |
| 16 | iv + Q | - | TRUE | TRUE | TRUE | TRUE | 48000 |
| 17 | I' + P | TRUE | - | - | - | - | 37142 |
| 18 | I' + Q | TRUE | - | - | - | TRUE | 36998 |
| 19 | II' + P | TRUE | - | - | TRUE | - | 40926 |
| 20 | II' + Q | TRUE | - | - | TRUE | TRUE | 40686 |
| 21 | III' + P | TRUE | - | TRUE | - | - | 37327 |
| 22 | III' + Q | TRUE | - | TRUE | - | TRUE | 37103 |
| 23 | IV' + P | TRUE | - | TRUE | TRUE | - | 45732 |
| 24 | IV' + Q | TRUE | - | TRUE | TRUE | TRUE | 45575 |
| 25 | i' + P | TRUE | TRUE | - | - | - | 40876 |
| 26 | i' + Q | TRUE | TRUE | - | - | TRUE | 40643 |
| 27 | ii' + P | TRUE | TRUE | - | TRUE | - | 42516 |
| 28 | ii' + Q | TRUE | TRUE | - | TRUE | TRUE | 42360 |
| 29 | iii' + P | TRUE | TRUE | TRUE | - | - | 44862 |
| 30 | iii' + Q | TRUE | TRUE | TRUE | - | TRUE | 44814 |
| 31 | iv' + P | TRUE | TRUE | TRUE | TRUE | - | 48855 |
| 32 | iv' + Q | TRUE | TRUE | TRUE | TRUE | TRUE | 48767 |

\* Formulacodes. For numerals with primes such as I' or ii', replace "SessionDay" with "Ltlog":
II: Generation + s(SessionDay)
III: s(SessionDay, by=Generation)
IV: s(SessionDay)
i: Generation \* SessionDay
ii: Generation + SessionDay
iii: Generation:SessionDay
iv: SessionDay
Lure attenuation pattern codes:
P: Company + s(Since.open, by=Company)

Models 1 and 2 were the standards used for the model family. For Model 1, temporal male population density fluctuation was smoothed by a spline and lure attractiveness was linearly attenuated on a log scale. For Model 2, pheromone attenuation was approximated by spline regression. They are expressed in R-style formula notation as follows:

~~~
M1 <-gam(N ∼ s(Site, bs=“re”) + Generation + s(SessionDay,
by=Generation) + Company + s(Since.open, by=Company), data=data,
family=nb(theta=NULL, link=“log”))
M2 <-gam(N ∼ s(Site, bs=“re”) + Generation + s(SessionDay,
by=Generation) + Company * Since.open, data=data,
family=nb(theta=NULL, link=“log”))
~~~

In the gam() function of the “mgcv” package of R, a random intercept was implemented with s(Variable, bs = “re”) where bs is the spline basis. The default setting for the smoother terms is s(bs = “tp”), which is fitted by a penalized thin-plate spline. Spline smoothness was automatically optimized.

The variables *τ* and *s*_*k,τ*_ were stored under “SessionDay” and “Since.open” in the dataset. The terms s(Session Day, by = Generation) and s(Since.open, by = Company) correspond to the smoothing terms *f*_*g*_(*τ*) and *ψ*_*j*_(*s*_*k,τ*_), respectively. Moth generation and lure manufacturer were included as covariates in the smoothing terms and stored as the data variables “Generation” and “Company,” respectively. The difference between Models 1 and 2 was the use of spline in the *s*_*k,τ*_ effect term. “Company * Since.open” in Model 2 corresponds to “Company + s(Since.open, by = Company)” in Model 1. Any smoothing term in “mgcv” was centered to mean = 0. Thus, Model 2 requires the separate “Company” term.

#### Models using light trap data as male density calibrator

Sixteen other candidate model comprising a model family based on Eq. 3 and Eq. 4 were constructed to evaluate lure attenuation after air exposure. Light trap data were used to calibrate daily male density. The Models 1 and 2 variants were Models 17 and 18, respectively (Table 2):

~~~
M17 <-gam(N ∼ s(Site, bs=“re”) + Generation + s(Ltlog,
by=Generation) + Company + s(Since.open, by=Company), data=data,
family=nb(theta=NULL, link=“log”))
M18 <-gam(N ∼ s(Site, bs=“re”) + Generation + s(Ltlog,
by=Generation) + Company * Since.open, data=data,
family=nb(theta=NULL, link=“log”))
~~~

where “Ltlog” corresponds to ln(*L*_g,r_ + 0.5) in Eq. 3.

#### Cross-validation of candidate models

Along with the aforementioned basic models, 32 candidate models were constructed and cross-validated. The generalization performance of the model was defined as the prediction accuracy for the daily capture on an unknown trap site. The models were fitted using count data from three out of four sites (A–D), in which the data for four insect generations were combined. The remaining unused site then served as the test data source.

The prediction accuracy was determined according to the negative log-likelihood (NLL) of *N*_*g,i,j,k,τ*_ observations under predictor size *μ*_*g,i,j,k,τ*_ in Eq. 4. A maximum of four test cases may be deployed. By excluding A, B, C, or D, NLL sums were obtained for four leave-one-out cross-validations. The model structure with the highest generalization performance (minimal NLL) was selected. For the best-fit model determined by the NLL values, the final parameters were fitted using full data for all four sites. The original dataset and the R code are provided as Supplementary Materials S1 and S2, respectively.

## 4. Results

During the 2017–2018 trap sessions, the temperature was lower for the 0^th^ (overwintering) *A. honmai* generation emerging in April than it was for the other moth generations that occurs in Shizuoka prefecture from late June to August (Table 1). Every generation was subjected to heavy rainfall (> 10 mm/day). The trap sessions covered nearly all 2^nd^ and 3^rd^ generation adult emergence periods in 2017. Trap settlement was delayed in the 1^st^ generation of 2017 and the 0^th^ generation of 2018 because pre-settlement lure exposure had to be planned retroactively based on long-term weather forecasts.

Using three pheromone traps, four sites, and 70 trap nights, we captured 8882, 9215, 17607, and 18728 *A. honmai* adults (median capture per trap per day = 4, 2, 7, and 8, respectively) in the 1^st^, 2^nd^, 3^rd^, and 0^th^ generations, respectively (Figure 2). The “SE” lures captured 29931 adult males or 5218, 6520, 10347, and 7846 in the 1^st^, 2^nd^, 3^rd^, and 0^th^ generations, respectively. The maximum and median captures per trap per day were 345 and 11, respectively. These rates were 1.8× and 3.8× larger than those obtained using the “SC” traps (16545 [2603, 1782, 4711, and 7449] individuals; daily maximum and median = 201 and 5, respectively) and “OT” traps (7956 [1061, 913, 2549, and 3433] individuals; daily maximum and median = 132 and 2, respectively).

**Fig 2.**
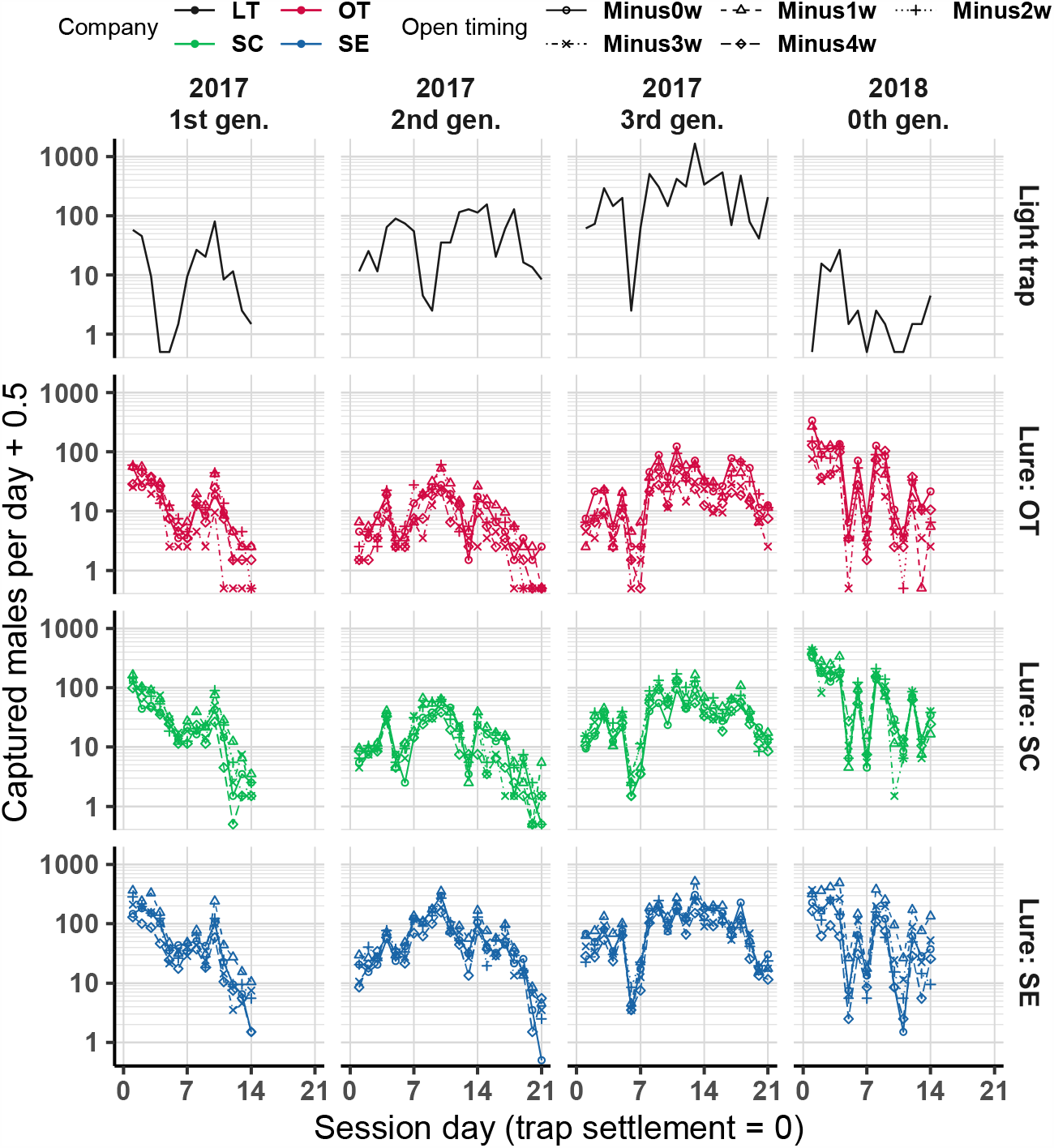
Raw capture during trap sessions. Numbers of captured males from four pheromone-trap sites are pooled.

In the light trap, we captured 269, 1164, 6347, and 64 adult males (median capture per trap per day = 9, 35, 206, and 1, respectively) in the 1^st^, 2^nd^, 3^rd^, and 0^th^ generations, respectively (Figure 2). Hence, *A. honmai* light trapping was less effective during the earlier seasons (0^th^ and 1^st^ generations) than it was in summertime.

### Cross validation of the candidate models

Of the 32 candidate models, Model 2 (total NLL = 32555) and Model 1 (total NLL = 32669) had the highest and second highest generalization performances, respectively (Table 2). Comparison of Models 1 and 2 suggested that linear regression may be appropriate for lure degradation after air exposure whereas base population dynamics must be fitted to nonlinear regression. The goodness-of-fit substantially declined after we assumed linearity for population density (Table 2: Model 1 *vs*. Model 9: *Δ*NLL = 7676; Model 2 *vs*. Model 10: *Δ*NLL = 7870).

### Models using pheromone traps with different lure ages

Based on Models 1 and 2, the final parameters were fitted with data from all four sites (Table 3). The AIC of Model 2 was 41.1 units higher than that of Model 1. The adjusted R^2^ were 0.36 and 0.326 for Models 1 and 2, respectively. Thus, the goodness-of-fit was slightly better for Model 1 (nonlinear attenuation) than Model 2 (linear attenuation).

**Table 3.** Summary table for GAMM assuming that lure attractiveness nonlinearly (Model 1) or linearly (Model 2) decreases after air exposure (“Since.open”). The reference levels for the parametric coefficients are “OT” and “0” for lure type (“Company”) and discrete moth generation per year (“Generation”), respectively. Smooth terms show daily fluctuations (“SessionDay”) in population abundance during sampling of each generation.

| Model variation | Model 1 (nonlinear attenuation) | Model 2 (linear attenuation) |
| --- | --- | --- |
| Parametric coefficients | Estimate (SE) | Estimate (SE) |
| Intercept | 135.2 (29.83) | 135.8 (29.79) |
| Generation = 1 | −134.8 (29.84) | −134.8 (29.80) |
| Generation = 2 | −134.5 (29.83) | −134.5 (29.79) |
| Generation = 3 | −133.7 (29.83) | −133.7 (29.79) |
| Company = SC | 0.7605 (0.03772) | 0.5126 (0.08489) |
| Company = SE | 1.513 (0.03685) | 1.334 (0.08303) |
| Since.open | — | −0.02564 (0.002597) |
| SC: Since.open | — | 0.01110 (0.003375) |
| SE: Since.open | — | 0.007713 (0.003295) |
| Smooth terms | edf (Chisq) | edf (Chisq) |
| s(Site) | 2.979 (426.73) | 2.979 (423.1) |
| s(SessionDay): Generation0 | 8.097 (745.32) | 8.094 (772.1) |
| s(SessionDay): Generation1 | 6.238 (772.70) | 6.238 (756.9) |
| s(SessionDay): Generation2 | 8.475 (708.74) | 8.474 (728.7) |
| s(SessionDay): Generation3 | 8.687 (797.49) | 8.683 (816.7) |
| s(Since.open): CompanyOT | 2.372 (98.98) | — |
| s(Since.open): CompanySC | 3.407 (69.22) | — |
| s(Since.open): CompanySE | 3.820 (75.77) | — |
| NB dispersion parameter $\theta$ | 1.457 | 1.434 |
| Deviance explained | 60.8% | 60.2% |
| Adjusted R<sup>2</sup> | 0.36 | 0.326 |
| AIC | 25037.16 | 25072.31 |

According to Model 2, the initial lure attractiveness values and rates of temporal lure attenuation differed among lure types. The standard lure attractiveness for this model was defined as the expected capture rate using the OT lure immediately after opening. For the SC lure, the expected capture immediately after opening was *e*^0.5126^ = 167% of OT. For the SE lure, it was *e*^1.334^ = 380% (Table 3). The attractiveness of all three lure types decreased with increasing number of days after opening (Figure 3b). The rate of attractiveness attenuation was commensurate with the size of the “Since.open” effect term in Model 2. The rate of attractiveness attenuation showed the largest decline in OT (−0.02564 ± 0.002597; estimated coefficient ± SE). The rates of attractiveness attenuation were −0.01454 and −0.01793 for the SC and SE lures, respectively. Daily lure attractiveness fell to 97.47% (OT), 98.56% (SC), and 98.22% (SE). These values were not derived from a single term. Rather, they were calculated as the sums of “Since.open” and “SC:Since.open” or “SE:Since.open.” hence, estimates of their intervals were not obtained for SC or SE.

**Fig 3.**
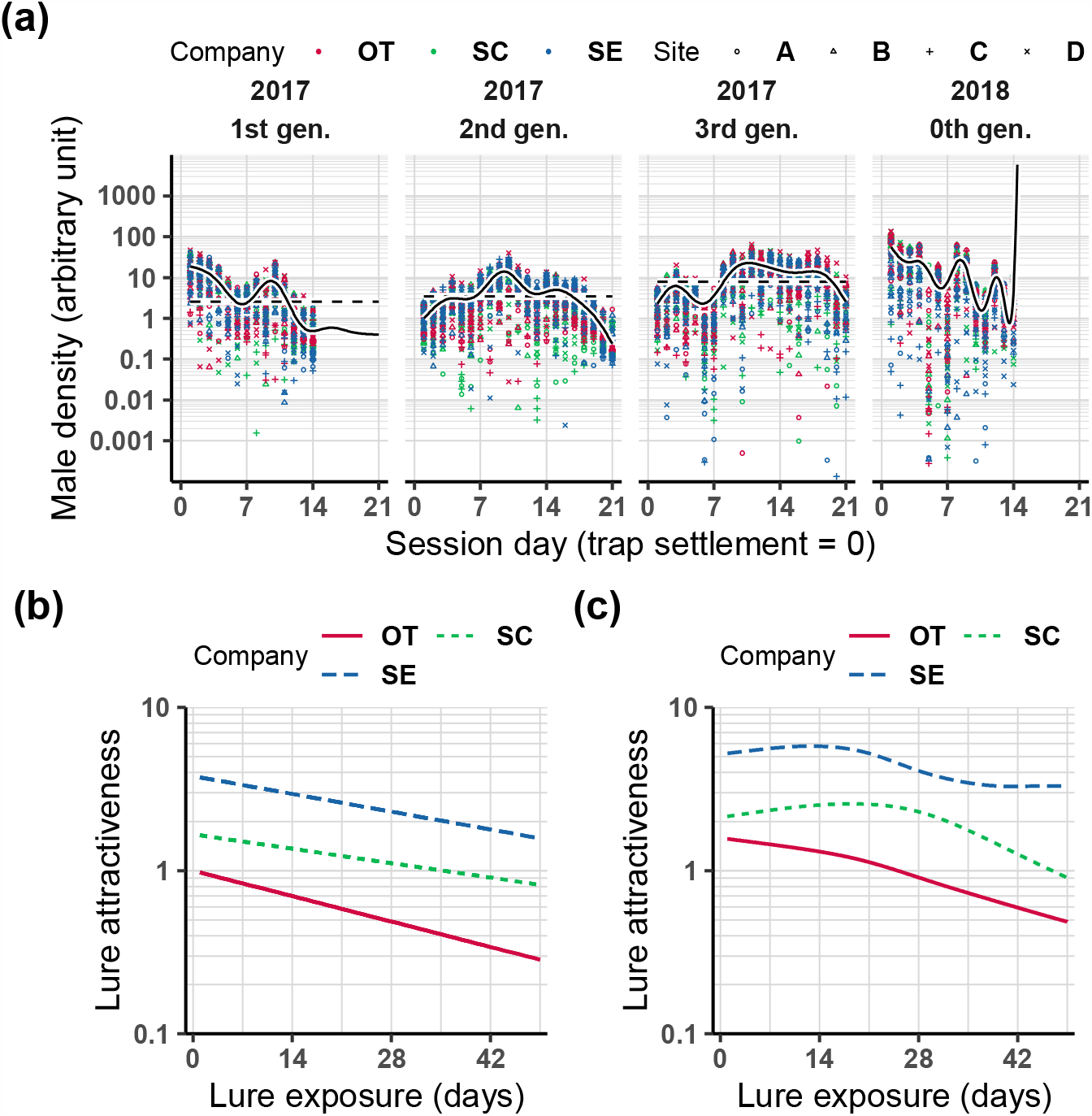
Graphical summary of GAMM Models 2 1 assuming exponential or nonlinear decrease in pheromone lure attractiveness, respectively. (a) Regional male density fluctuation was estimated as sum of effect terms “intercept + Generation + s(SessionDay, by = Generation).” Model 2 is plotted here and Model 1 was nearly identical to it. Dots signify partial residuals for each data point. Horizontal broken lines represent size of “intercept + Generation.” This parameter is not shown for Generation 0 as the intercepts were estimated beyond the plot range because of extrapolation of the periods after day 14. (b) Fluctuations in lure product attractiveness were predicted from Model 2 and corresponded to the effect terms “Company * Since.open.” (c) Lure attractiveness was predicted from Model 1 and corresponding to “s(Since.open, by = Company).”

### Models using light trap data as male density calibrator

The model family using light trap data as a predictor of population fluctuation performed poorly (Table 2). The best in this family was Model 18 (sum NLL = 36998, adjusted R^2^ = 0.332) assuming “Generation + s(Ltlog, by = Generation)” and “Company * Since.open” for base density and lure attenuation, respectively. Model 18 was the counterpart of Model 2. The second best was Model 22 (37103, adjusted R^2^ = 0.33) which was a Model 18 variant because its base density term was altered to “s(Ltlog, by = Generation).” The third best was Model 17 (sum NLL = 37142, adjusted R^2^ = 0.34) which was also a Model 18 variant as its lure attenuation term was changed to the nonlinear term “Company + s(Ltlog, by = Company).” Model 17 was the counterpart of Model 1.

The aforementioned models all had the “s(…, by = Generation)” covariate whereas others such as Model 20 (sum NLL = 40686) with relatively poor performance lacked that term. Thus, no single regression curve or line could convert all light to pheromone trap capture data or vice-versa. Therefore, the data must be individually converted for each generation via nonlinear regression such as spline (Figure 4a; four regression curves per generation).

**Fig 4.**
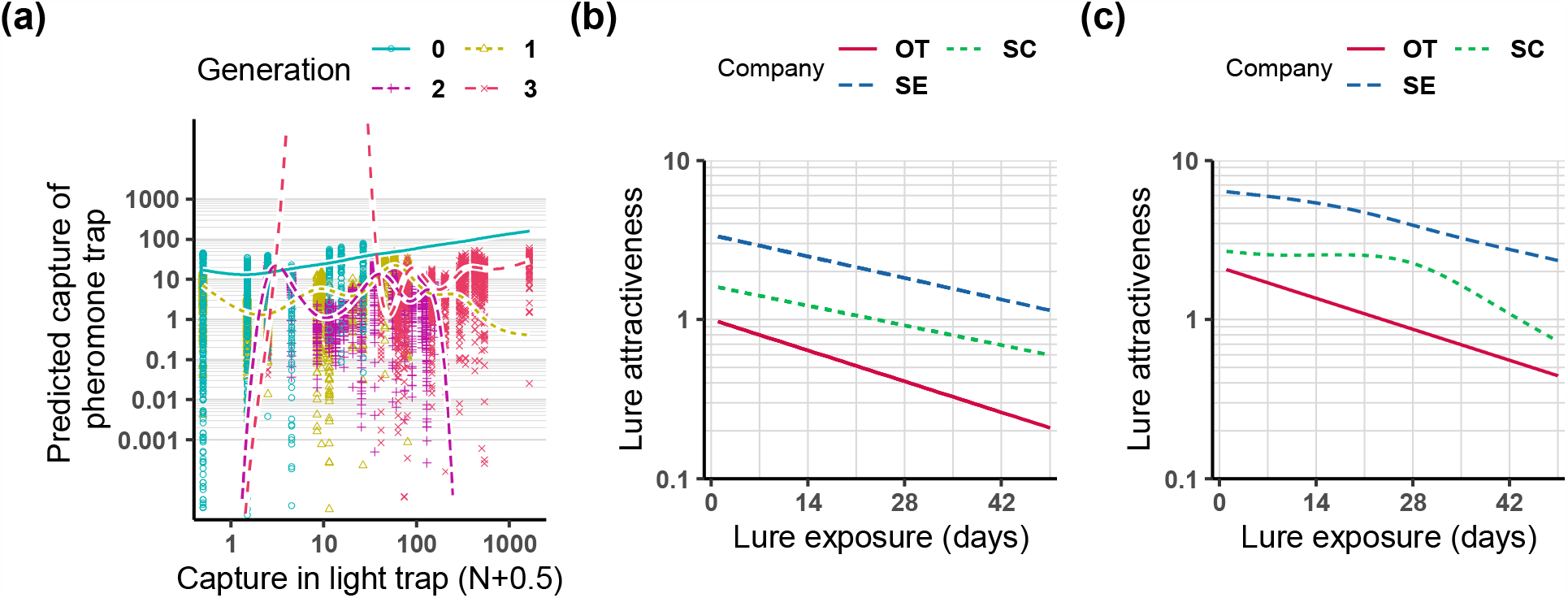
Graphical summary of GAMM structures using light trap data. (a) Performance of light trap data (Ltlog: x-axis) as predictor of capture with pheromone lure on same day (y-axis). Regression curves are defined as sum of effect terms “intercept + Generation + s(Ltlog, by = Generation).” Points signify partial residual. Though plots only show result of Model 18 (assuming exponential decrease in pheromone lure attractiveness), Model 17 (assuming nonlinear decrease in pheromone lure attractiveness) had nearly same result. (b) Fluctuations in lure product attractiveness were predicted from Models 18 and (c) 17. Both are plotted in same way as Models 2 and 1, respectively.

Generalization performance was comparatively lower for models using light trap data. Nevertheless, the lure attenuation patterns estimated by Models 18 (Figure 4b) and 17 (Figure 4c) resembled those estimated by Models 2 (Figure 3b) and 1 (Figure 3c), respectively. If we had adopted Model 18 as the final model, the initial attractiveness efficacies of the SC-and SE-type lures would be 162% and 340%, respectively, relative to OT. For the lure attenuation rate, the OT slope was −0.03200 ± 0.00278 (estimate ± SE) while those for the SC and SE lures were −0.0204 and −0.0223, respectively. Hence, daily lure attractiveness declined to 96.85%, 97.98%, and 97.80% for OT, SC, and SE, respectively.

## 5. Discussion

Previous studies used nonlinear regression methods such as GAM to segregate background insect population density from long-term insect population monitoring data (Benton, Bryant, Cole, & Crick, 2002; Klapwijk, Csóka, Hirka, & Björkman, 2013; Yamamura, 2016). Here, we used GAMM to quantify the temporal dynamics of the attractiveness of tortrix moth pheromone lures separately from fluctuations in insect population density. We combined lures with different opening (air exposure) timings and used them as an internal reference to correlate the effects of lure age with the observed insect number.

In the supported model (Table 3: Model 2), the base dynamics were fitted by a nonlinear regression. In contrast, linear regression was better suited to fit lure attenuation after field settlement. The nonlinear regression (Table 3: Model 1) showed slightly better goodness-of-fit (adjusted R^2^) and AIC than the linear regression. Nevertheless, the sum negative log likelihood (NLL) for the unknown trap sites suggest that linear regression of lure attenuation would be more practical than nonlinear regression in terms of generalization performance and ease of data conversion.

Pest insects are often monitored with pheromone traps to detect peak regional adult emergence timing, or evaluate local population abundance. In both cases, unbiased indices of regional density must be determined for each time point. In our model structure, *d*_*g,i,τ*_ corresponded to fluctuations in regional male density. For *A. honmai*, however, the expected male moth capture rate exponentially decreased with lure age. Therefore, the male density estimate would be biased if we did not eliminate the effect of lure attenuation. If we allow the lure attractiveness to decline to 90% of its initial value, then Model 2 would disclose that lure attenuation is non-negligible at 4.11 d, 7.25d, and 5.88 d after settlement of the OT, SC, and SE lures, respectively.

GAMM enables us to utilize trap data for relatively longer periods as it calibrates heterogeneous trap records with various lure types and conditions. An intuitive density index is equivalent to *d*_*g*_ + *f*_*g*_(*τ*) (mean generation density + temporal fluctuation) (Figure 3a). In biological terms, it represents the expected daily number of males captured by a trap equipped with a brand-new lure (note that this value is not an absolute density such as individuals/m^2^). If we adhere to the R mgcv package, we first calculate the predicted values via the “predict(M2, type=‘terms’)” command; then the estimate is given as the sum of the effects “Intercept + Generation + s(SessionDay, by = Generation)” (also see ESM S2).

While *d*_*g,i,τ*_ corresponds to male density, *ψ*_*j*_(*s*) corresponds to the time course of pheromone lure attractiveness. The latter was determined from the evaporation of the pheromone components from the lure dispenser. A classical first-order model has been used to explain the measured exponential decrease in evaporation rate (Heuskin et al., 2011; McDonough & Butler, 1983; McDonough, Aller, & Knight, 1992). In a first-order relationship, the release rates are proportional to the volatile content in the dispenser. Both the volatile content and the evaporation rates linearly decrease on a logarithmic scale. For tea tortricid moths, Tamaki & Noguchi (1983) used this assumption to estimate the rates of decline in the pheromone components for *A. honmai* and its co-occurring species *Homona magnanima* Diakonoff (Tortricidae). They prepared synthetic pheromone by mixing Z9-14Ac, Z11-14Ac, E11-14Ac, and 10-Me-12Ac at 70:30:10:200 (23%, 10%, 3%, and 64%). This formulation resembled that of the OT lure. In the dispenser, its half-lives were ∼10 and ∼30 days when the volatiles were loaded onto a plastic cap or a rubber septum, respectively.

Tamaki & Noguchi (1983) also found that the half-life of the E11-14Ac content was nearly twice that of the major two components, Z9-14Ac and Z11-14Ac, whereas the half-life of 10-Me-12Ac was half that of the two especially when they were loaded on a rubber septum. However, the authors argued that the heterogeneous pheromone evaporation rates did not markedly affect lure attractiveness stability. The minor components had wider ranges at the optimal mixing ratios than the major two. Pheromonal activity persisted in *H. magnanima* as long as Z9-14Ac:Z11-14Ac was kept 3:1 (Noguchi et al., 1981). According to Tamaki & Noguchi (1983), the four components were kept for 30 days and 60 days within the optimal mixing ratio for *A. honmai* males on a plastic cap or a rubber septa, respectively. They concluded that the exhaust of the volatiles, rather than the deviation from the optimal mixing ratio, were responsible for the persistence.

Regarding OT lure trap data, attractiveness could be calibrated based on our Model 2 as this pheromone lure decreases rapidly but shows strong linearity. In contrast, according to Model 1 (nonlinear version), the attractiveness of the SC and SE lures slightly decreased initially but rapidly declined thereafter (Figure 3c). Therefore, uncalibrated capture data could be used for SC and SE. In this case, we should truncate data gathered > 1 month after the pheromone lure settlement dates just as survey guidelines have recommended (Tatara & Kodomari, 2000).

The first-order assumption clearly explains the relationship between exposure time and pheromone evaporation rate (≈ dose). However, another assumption is required to explain our result that the *A. honmai* male captures decreased exponentially after unsealing the lures, especially the OT lures. Although there has been a long-standing debate how should we model the diffusion of pheromone plume in the ambient, studies have assumed the area of the attraction range (∝ expected capture if the males distribute uniformly) increases linearly first with the pheromone dose (Branco, Jactel, Franco, & Mendel, 2006; Turchin & Odendaal, 1996; Wall & Perry, 1987). However, there seems a saturation at higher dose range; the cause has been considered the limit in the flyable distance of the male insects (Östrand & Anderbrant, 2003; Wall & Perry, 1987). Regarding *A. honmai*, table 7 of Tamaki et al. (1980) shows that male *A. honmai* capture is initially linearly correlated with synthetic pheromone dose (at a 63:31:4:200 [21%, 10%, 1%, and 68%] mixing ratio) approximately on a log-log scale. Thence, capture reaches a plateau as the initial load increases beyond 10 mg/dispenser.

These properties of *A. honmai* pheromone could explain why the attenuation of SC/SE lure attractiveness was relatively slow and nonlinear. The threshold level reported by Tamaki et al. (1980) approached the initial loads of OT and SC. Therefore, the commercial lure yields may also fluctuate around the threshold immediately after lure settlement. Thereafter, the expected capture started to decline exponentially. For OT, the dispenser substrate was a plastic capsule whereas it was a rubber septum for SC and SE. The latter provided a pheromone dose half-life that was threefold longer than those of the former. Overall, the pheromone charge in the OT lure underwent exponential decrease very soon after opening whereas the SC and SE dose levels remained on a plateau for the first month after opening.

Caution is needed when interpreting the estimated parameter values using datasets for multiple trap sessions with different time periods. For Model 2, the intercept was ∼135.8 (Table 3), which was not a plausible density range of insects. It was generated when the mgcv::gam() function tried to centralize the smoother term s(SessionDay, by = Generation) for the entire [1, 21] domain over all four generations. Nevertheless, the actual data only covered [1, 14] for the 1^st^ and 0^th^ generations (Figure 3a) and extrapolation occurred.

The extrapolation may be avoided by writing and running the models with probabilistic programming languages such as JAGS or Stan. However, the GAM(M) approach is more reproducible and feasible for agronomists and policymakers. In GAMM, the intercept shift was compensated by other terms. We can nonetheless predict regional male density in the GAMM using the sum of “intercept + Generation + s(SessionDay, by = Generation)” (Figure 3a). As it is expressed as the sum of multiple terms, we are unable to obtain interval estimates such as prediction intervals for male density. Though this problem persists with GAM(M), we may obtain interval estimates by sampling from posterior distributions when the model structure is ported to the Monte Carlo methods.

Our model comparison demonstrated no universal correlation between pheromone and light trap records for the same day. The light trap was relatively more efficient in the summertime (generations with high population densities) but not during earlier seasons. Similar trends were reported for other lepidopteran (Campbell, Walgenbach, & Kennedy, 1992; Delisle, West, & Bowers, 1998; Tanaka, Kouno, Hirose, & Futai, 1995) and hemipteran (Nakamura & Nishino, 1999) pest species. The mechanism might entail interference between pheromone lure attractiveness and female moths, which leads to decreased lure attractiveness at higher densities. However, this theory has not yet been fully corroborated by field experiments (Campbell, Walgenbach, & Kennedy, 1992; Laurent & Frérot, 2007).

As there is only a weak correlation between the light trap and pheromone trap capture records (Figure 4a), straightforward data conversion between them is improbable. As relative density estimates are saturated by method-specific constraints, we require an absolute density measurement to integrate heterogeneous monitoring data and investigate the dynamics of long-term population fluctuations. Absolute density estimation with pheromone traps has been conducted via marked insect release-recapture (Adams et al., 2017; Arakaki et al., 2008; Östrand & Anderbrant 2003; Turchin & Odendaal, 1996). Going forward, our model structure will be updated to enable inference on the absolute density. Such an update includes a more full-fledged state-space model framework because the research design of release-recapture requires the implementation of intrinsic population dynamics such as adult emergence, survival, and possibly the frequency and distance of flight as well as the mating motivation, which may also vary during the lifespan of adult males.

## Supporting information

ESM1 (raw data) and ESM2 (R script)

## Data availability

The data source used in this study is archived on ‘https://doi.org/10.6084/m9.figshare.13206236.v1‘.

## Author contribution

SY designed the field experiments and collected the pheromone trap data. SY and YH collected the light trap data. SM created the statistical framework and wrote the first draft. All authors wrote the final manuscript.

## Table and figure captions

Supplementary Materials S1

Dataset: Captured males

Supplementary Materials S2

R source code

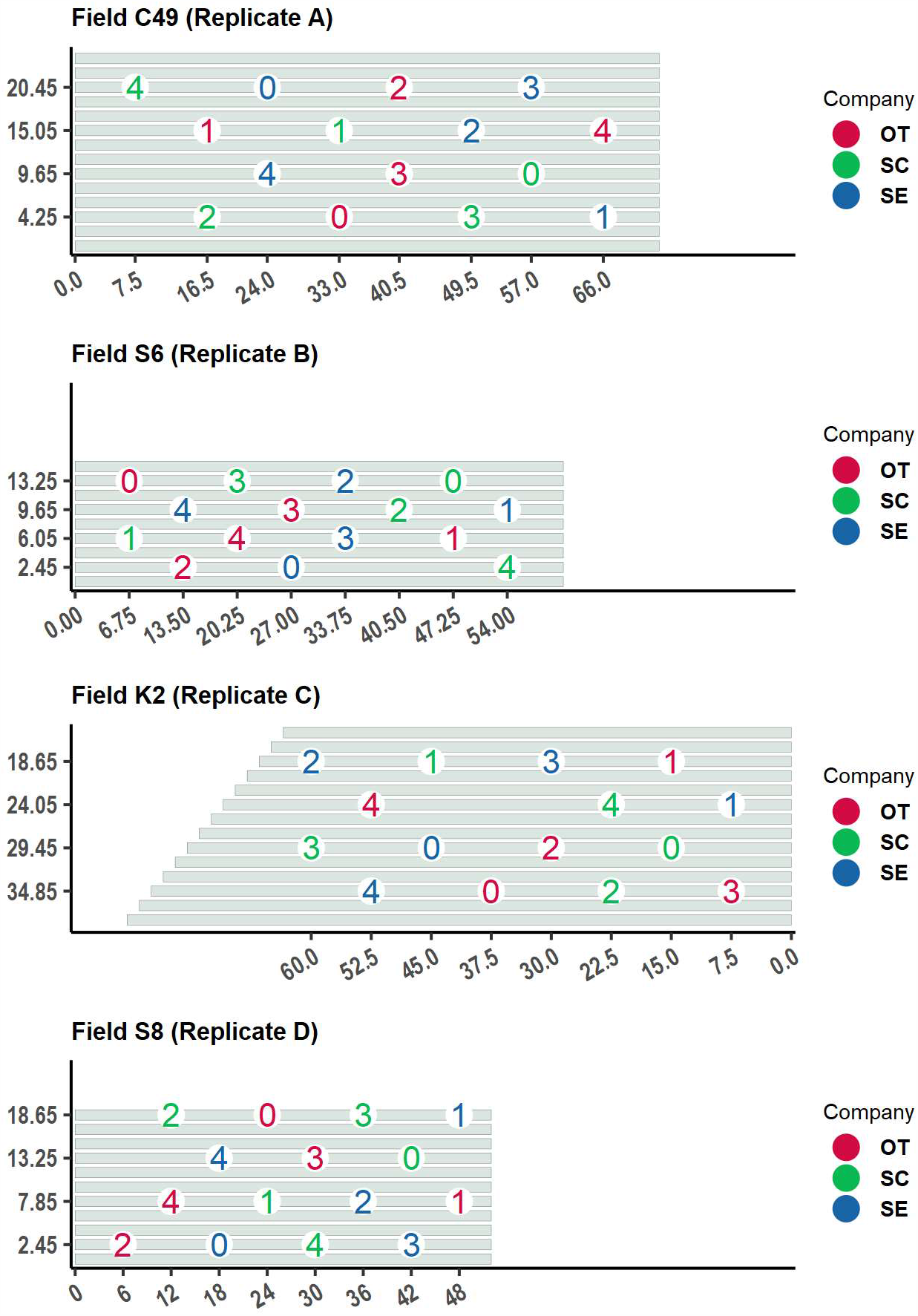
Location of traps in each tea field shown in Figure 1. Right direction points north. Scale in meters indicates distances from ends of tea hedges. Each hedge is 1.3 m wide and each furrow is 50 cm wide. Numbers on plot represent weeks of pre-settlement lure exposure.

## References

Adams, C. G., Schenker, J. H., McGhee, P. S., Gut, L. J. Brunner, J. F., & Miller, J. R. (2017). Maximizing information yield from pheromone-baited monitoring traps: estimating plume reach, trapping radius, and absolute density of Cydia pomonella (Lepidoptera: Tortricidae) in Michigan apple. Journal of Economic Entomology, 110, 305–318. doi: 10.1093/jee/tow258

Agresti, A. (2002). Categorical data analysis. 2nd ed. Hoboken, NJ: John Wiley & Sons.

Arakaki, N., Nagayama, A., Kobayashi, A., et al. (2008). Estimation of abundance and dispersal distance of the sugarcane click beetle Melanotus sakishimensis Ohira (Coleoptera: Elateridae) on Kurima Island, Okinawa, by mark-recapture experiments. Applied Entomology and Zoology, 43, 409–419. doi: 10.1303/aez.2008.409

Benton, T. G., Bryant, D. M., Cole, L., & Crick, H. Q. (2002). Linking agricultural practice to insect and bird populations: a historical study over three decades. Journal of Applied Ecology, 39, 673–687. doi: 10.1046/j.1365-2664.2002.00745.x

Bonebrake, T. C., Christensen, J., Boggs, C. L., & Ehrlich, P. R. (2010). Population decline assessment, historical baselines, and conservation. Conservation Letters, 3, 371–378. doi: 10.1111/j.1755-263X.2010.00139.x

Branco, M., Jactel, H., Franco, J. C., & Mendel, Z. (2006). Modelling response of insect trap captures to pheromone dose. Ecological Modelling, 197, 247–257. doi: 10.1016/j.ecolmodel.2006.03.004

Butler L. I., & McDonough, L. M. (1979). Insect sex pheromones. Evaporation rates of acetates from natural rubber septa. Journal of Chemical Ecology, 5, 825–837. doi: 10.1007/BF00986567

Campbell, C. D., Walgenbach, J. F., & Kennedy, G. G. (1992). Comparison of black light and pheromone traps for monitoring Helicoverpa zea (Boddie) (Lepidoptera: Noctuidae) in tomato. Journal of Agricultural Entomology, 9, 17–24.

Cardé, R. T. (1976). Utilization of pheromones in the population management of moth pests. Environmental Health Perspectives, 14, 133–144. doi: 10.2307/3428371

Delisle, J., West, R. J., & Bowers, W. W. (1998). The relative performance of pheromone and light traps in monitoring the seasonal activity of both sexes of the eastern hemlock looper, Lambdina fiscellaria fiscellaria. Entomologia Experimentalis et Applicata 89, 87–98. doi: 10.1046/j.1570-7458.1998.00385.x

Elton, C., & Nicholson, M. (1942). The ten-year cycle in numbers of the lynx in Canada. Journal of Animal Ecology, 11, 215–244. doi: 10.2307/1358

Fukaya, K. (2016). Time series analysis by state space models and its application in ecology. Japanese Journal of Ecology, 66, 375–389. doi: 10.18960/seitai.66.2_375

Heuskin, S., Verheggen, F. J., Haubruge, E., Wathelet, J.-P., & Lognay, G. (2011). The use of semiochemical slow-release devices in integrated pest management strategies. BASE, 11, 459–470.

Ishijima, C., Sato, Y., & Ohtaishi, M. (2009). Occurrence of the tea tortrix complex and their parasitoids in pesticide-free tea fields in Shizuoka Prefecture, Japan. Tea Research Journal, 2009, 108–118. doi: 10.5979/cha.2009.108_7

Japan Plant Protection Association. (2000). Guidebook for forecasting and control of pests use pheromones. Tokyo: Japan Plant Protection Association.

Jorgensen, C., Enberg, K., Dunlop, E.S., et al. (2007). Ecology: Managing evolving fish stocks. Science, 318, 1247–1248. doi: 10.1126/science.1148089

Kawasaki, K., Nakamura, K., Noguchi, H., Sugie, H., & Tamaki, Y. (1979). 323 Effective range and time of attraction of the smaller tea tortrix moth to sex pheromone trap (チャノコカクモンハマキ性フェロモントラップの有効範囲および誘引時刻). Abstract of the 23th Annual Meeting of Japanese Society of Applied Entomology and Zoology (p. 78).

Kawasaki, K., & Tamaki, Y. (1980). Relationship between sex pheromone trap position and capture of adult males of Adoxophyes sp. Japanese Journal of Applied Entomology and Zoology, 24, 253–255. doi: 10.1303/jjaez.24.253

Kéry, M., & Schaub, M. (2011). Bayesian population analysis using WinBUGS: a hierarchical perspective. London: Academic Press.

Klapwijk, M. J., Csóka, G., Hirka, A., & Björkman, C. (2013). Forest insects and climate change: long-term trends in herbivore damage. Ecology and Evolution, 3, 4183–4196. doi: 10.1002/ece3.717

Laurent, P., & Frérot, B. (2007). Monitoring of European corn borer with pheromone-baited traps: review of trapping system basics and remaining problems. Journal of Economic Entomology, 100, 1797–1807. doi: 10.1603/0022-0493(2007)100[1797:moecbw]2.0.co;2

McDonough, L. M., Aller, W. C., & Knight, A. L. (1992). Performance characteristics of a commercial controlled-release dispenser of sex pheromone for control of codling moth (Cydia pomonella) by mating disruption. Journal of Chemical Ecology, 18, 2177–2189. doi: 10.1007/BF00984945

McDonough, L. M., & Butler, L. I. (1983). Insect sex pheromones: Determination of half-lives from formulations by collection of emitted vapor. Journal of Chemical Ecology, 9, 1491–1502. doi: 10.1007/BF00988515

Nabeta, F. H., Nakai. M., & Kunimi, Y. (2005). Effects of temperature and photoperiod on the development and reproduction of Adoxophyes honmai (Lepidoptera: Tortricidae). Applied Entomology and Zoology, 40, 231–238.

Nakamura, Y., Nishino, T. (1999). Forecasting fruit damage caused by the brown-winged green bug, Plautia stali Scott, and its occurrence using synthetic aggregation pheromone traps. Kyushu Plant Protection Research, 45, 119–122.

Noguchi, H., Tamaki, Y., Arai, S., Shimoda, M., & Ishikawa, I. (1981). Field evaluation of synthetic sex pheromone of the Oriental Tea Tortrix Moth, Homona magnanima Diakonoff (Lepidoptera: Tortricidae). Japanese Journal of Applied Entomology and Zoology, 25, 170–175. doi: 10.1303/jjaez.25.170

Oba, M. (1979). Seasonal occurrence of the smaller tea tortrix moth, Adoxophyes sp., as measured with a pheromone trap in the tea field. Tea Research Journal, 1979, 6–11. doi: 10.5979/cha.1979.50_6

Osada, Y., Yamakita, T., Shoda-Kagaya, E., Liebhold, A. M., & Yamanaka, T. (2018). Disentangling the drivers of invasion spread in a vector-borne tree disease. Journal of Animal Ecology, 87, 1512–1524. doi: 10.1111/1365-2656.12884

Osakabe, M. (1985). Yearly changes of the number of tea insect pest adults came flying to a blueness fluorescent light trap during 1947 to 1979. Tea Research Journal, 1985, 40–45. doi: 10.5979/cha.1985.62_40

Osakabe, M. (1986). Seasonal change of the number of tea insect pest adult came flying to a light trap in the tea field of our research institute. Tea Research Journal, 1986, 11–19. doi: 10.5979/cha.1986.11

Östrand, F., & Anderbrant, O. (2003). From where are insects recruited? A new model to interpret catches of attractive traps. Agricultural and Forest Entomology, 5, 163–171. doi: 10.1046/j.1461-9563.2003.00174.x

R Core Team. (2019). R version 3.6.1. https://www.R-project.org/

Reinke, B. A., Miller, D. A., & Janzen, F. J. (2019). What have long-term field studies taught us about population dynamics? Annual Review of Ecology, Evolution, and Systematics, 50, 261–278. doi: 10.1146/annurev-ecolsys-110218-024717

Royama, T. (1996). Analytical population dynamics, Dordrecht: Springer Science & Business Media. doi: 10.1007/978-94-011-2916-9

Sato, Y., Takeda, M., Onoda, H., & Takaoka, H. (2005). Practicality of automatic recording electrocution pheromone trap “moth-Counter” in forecasting of the smaller tea tortrix, Adoxophyes honmai Yasuda. Tea Research Journal, 21–29. doi: 10.5979/cha.2005.21

Sudo, M., Yamanaka, T., & Miyai, S. (2019). Quantifying pesticide efficacy from multiple field trials. Population Ecology, 61, 450–456. doi: 10.1002/1438-390X.12019

Tamaki, Y., Noguchi, H. (1983). Decrease rates of pheromonal components in attractant-formulations for tea tortricid moths under field conditions. Japanese Journal of Applied Entomology and Zoology, 27, 154–156. doi: 10.1303/jjaez.27.154

Tamaki, Y., Noguchi, H., Sugie, H., Sato, R, & Kariya, A. (1979). Minor components of the female sex-attractant pheromone of the smaller tea tortrix moth (Lepidoptera: Tortricidae): Isolation and identification. Applied Entomology and Zoology, 14, 101–113. doi: 10.1303/aez.14.101

Tamaki, Y., Noguchi, H., Sugie, H., Kariya, A., Arai, S., Ohba, M., Terada, T., Suguro, T., & Mori, K. (1980). Four-component synthetic sex pheromone of the smaller tea tortrix moth: field evaluation of its potency as an attractant for male moth. Japanese Journal of Applied Entomology and Zoology, 24, 221–228. doi: 10.1303/jjaez.24.221

Tamaki, Y., Noguchi, H., Yushima, T., & Hirano, C. (1971) Two sex pheromones of the smaller tea tortrix: Isolation, identification, and synthesis. Applied Entomology and Zoology, 6, 139–141. doi: 10.1303/aez.6.139

Tanaka, H., Kouno, S., Hirose, T., & Futai, K. (1995) Seasonal occurrence of lawn grass pests by light trap and pheromone trap on the turf ground in Hyogo Prefecture. Bulletin of the Hyogo Prefectural Agricultural Institute. Agricultural section, 43, 55–60.

Tatara, A., & Kodomari, S. (2000). Adoxophyes honmai (tea). In: Guidebook for Forecasting and Control of Pests Use Pheromones. Tokyo: Japan Plant Protection Association. pp 30–32.

Turchin, P., & Odendaal, F. J. (1996). Measuring the effective sampling area of a pheromone trap for monitoring population density of southern pine beetle (Coleoptera: Scolytidae). Environmental Entomology, 25, 582–588. doi: 10.1093/EE/25.3.582

Wall, C., & Perry, J. N. (1987). Range of action of moth sex-attractant sources. Entomologia Experimentalis et Applicata, 44, 5–14. doi: 10.1111/j.1570-7458.1987.tb02232.x

Witzgall, P., Kirsch, P., & Cork, A. (2010). Sex pheromones and their impact on pest management. Journal of Chemical Ecology, 36, 80–100. doi: 10.1007/s10886-009-9737-y

Wood, S. (2020). mgcv: Mixed GAM computation vehicle with automatic smoothness estimation. Version 1.8-33. https://CRAN.R-project.org/package=mgcv

Yamamura, K. (2016). Applying state-space models to the dynamics of insect populations. Japanese Journal of Ecology, 66, 339–350. doi: 10.18960/seitai.66.2_339

Yamamura, K. (1999). Transformation using (x+ 0.5) to stabilize the variance of populations. Population Ecology, 41, 229–234. doi: 10.1007/s101440050026

Yamamura, K., Yokozawa, M., Nishimori, M., Ueda, Y, & Yokosuka, T. (2006). How to analyze long-term insect population dynamics under climate change: 50-year data of three insect pests in paddy fields. Population Ecology, 48, 31–48. doi: 10.1007/s10144-005-0239-7

Yamanaka, T., Nelson, W. A., Uchimura, K., & Bjørnstad, O. N. (2012). Generation separation in simple structured life cycles: Models and 48 years of field data on a tea tortrix moth. American Naturalist, 179, 95–109. doi: 10.1086/663201

Yoshioka, T., & Sakaida, T. (2008). Control effect of communication disruption using synthetic sex pheromones for the smaller tea tortrix, Adoxophyes honmai (Yasuda) (Lepidoptera: Tortricidae), in sloped tea fields. Bulletin of the Fukuoka Agricultural Research Center, 27, 111–115.

