## Supplementary material for "Time-course in attractiveness of pheromone lure on the smaller tea tortrix moth: a generalized additive mixed model approach": ESM1 (raw data) and ESM2 (R script): S2_code.R

```
################################################################################################################################
Time-course in attractiveness of pheromone lure on the smaller tea tortrix moth: A generalized additive mixed model approach

Masaaki Sudo*1, Yasushi Sato1, Hiroshi Yorozuya1

1 Tea Pest Management Unit, Institute of Fruit Tree and Tea Science, NARO: Kanaya Tea Research Station, 2769, Shishidoi, Kanaya, Shimada, Shizuoka 428-8501, Japan

* To whom correspondence
ORCID ID:
0000-0001-9834-9857 (Masaaki Sudo)


> sessionInfo()
R version 3.6.1 (2019-07-05)
Platform: x86_64-w64-mingw32/x64 (64-bit)
Running under: Windows 10 x64 (build 19042)

Matrix products: default

Random number generation:
RNG:     Mersenne-Twister
Normal:  Inversion
Sample:  Rounding

locale:
[1] LC_COLLATE=Japanese_Japan.932  LC_CTYPE=Japanese_Japan.932    LC_MONETARY=Japanese_Japan.932
[4] LC_NUMERIC=C                   LC_TIME=Japanese_Japan.932

attached base packages:
[1] stats     graphics  grDevices utils     datasets  methods   base

other attached packages:
[1] gratia_0.4.1    devEMF_4.0      patchwork_1.0.1 ggthemes_4.2.0  extrafont_0.17  mgcv_1.8-33     nlme_3.1-149
[8] lubridate_1.7.9 magrittr_1.5    forcats_0.5.0   stringr_1.4.0   dplyr_1.0.0     purrr_0.3.4     readr_1.3.1
[15] tidyr_1.1.0     tibble_3.0.3    ggplot2_3.3.2   tidyverse_1.3.0

loaded via a namespace (and not attached):
[1] tidyselect_1.1.0 splines_3.6.1    haven_2.3.1      lattice_0.20-41  colorspace_1.4-2 vctrs_0.3.4
[7] generics_0.0.2   utf8_1.1.4       blob_1.2.1       rlang_0.4.7      pillar_1.4.6     glue_1.4.2
[13] withr_2.2.0      DBI_1.1.0        dbplyr_1.4.4     modelr_0.1.8     readxl_1.3.1     lifecycle_0.2.0
[19] munsell_0.5.0    gtable_0.3.0     cellranger_1.1.0 rvest_0.3.6      mvnfast_0.2.5    fansi_0.4.1
[25] Rttf2pt1_1.3.8   broom_0.7.0      Rcpp_1.0.5       scales_1.1.1     backports_1.1.9  jsonlite_1.7.0
[31] farver_2.0.3     fs_1.4.2         digest_0.6.25    hms_0.5.3        stringi_1.4.6    cowplot_1.0.0
[37] grid_3.6.1       cli_2.0.2        tools_3.6.1      crayon_1.3.4     extrafontdb_1.0  pkgconfig_2.0.3
[43] ellipsis_0.3.1   Matrix_1.2-18    xml2_1.3.2       reprex_0.3.0     assertthat_0.2.1 httr_1.4.2
[49] rstudioapi_0.11  R6_2.4.1         compiler_3.6.1

################################################################################################################################


# Preparation
library(tidyverse)
library(magrittr)
library(lubridate)
library(mgcv)

lure.sub <- readr::read_csv("S1_data.csv", col_types="DiiicciciiDDi")
summary(lure.sub)
lure.sub
xtabs(~Open.str+Generation, data=lure.sub)

dat <- lure.sub %>%
    dplyr::mutate(  Site=as.factor(Site),
                    Generation=as.factor(Generation),
                    Company=as.factor(Company),
                    Ltlog=log(N.lt+0.5)  ) %>%
    data.frame()
head(dat)

lure.allsites <- lure.sub %>%
    dplyr::group_by(Year, Generation, Open.str, Company, SessionDay) %>%
    tidyr::nest() %>%
    dplyr::mutate(N4=purrr::map_int(.x=data, .f=~sum(pluck(., "N")))) # sum of four sites
lure.allsites

lure.lt <- lure.sub %>%
    dplyr::group_by(Year, Generation, SessionDay) %>%
    tidyr::nest() %>%
    dplyr::mutate(N.lt=purrr::map_dbl(.x=data, .f=~median(pluck(., "N.lt")))) %>%
    dplyr::mutate(Company="LT")
lure.lt


> lure.sub
# A tibble: 4,200 x 13
   Date.end    N.lt  Year Generation Site  Company  Open Open.str SessionDay Since.open Date.open  Date.start     N
   <date>     <int> <int>      <int> <chr> <chr>   <int> <chr>         <int>      <int> <date>     <date>     <int>
 1 2017-06-27    57  2017          1 A     SC          0 Minus0w           1          1 2017-06-26 2017-06-26    69
 2 2017-06-27    57  2017          1 B     SC          0 Minus0w           1          1 2017-06-26 2017-06-26    23
 3 2017-06-27    57  2017          1 C     SC          0 Minus0w           1          1 2017-06-26 2017-06-26    27
 4 2017-06-27    57  2017          1 D     SC          0 Minus0w           1          1 2017-06-26 2017-06-26    15
 5 2017-06-27    57  2017          1 A     SC          1 Minus1w           1          8 2017-06-19 2017-06-26    25
 6 2017-06-27    57  2017          1 B     SC          1 Minus1w           1          8 2017-06-19 2017-06-26    61
 7 2017-06-27    57  2017          1 C     SC          1 Minus1w           1          8 2017-06-19 2017-06-26    71
 8 2017-06-27    57  2017          1 D     SC          1 Minus1w           1          8 2017-06-19 2017-06-26     8
 9 2017-06-27    57  2017          1 A     SC          2 Minus2w           1         15 2017-06-12 2017-06-26    54
10 2017-06-27    57  2017          1 B     SC          2 Minus2w           1         15 2017-06-12 2017-06-26    19
# ... with 4,190 more rows

> xtabs(~Open.str+Generation, data=lure.sub)
         Generation
Open.str    0   1   2   3
  Minus0w 168 168 252 252
  Minus1w 168 168 252 252
  Minus2w 168 168 252 252
  Minus3w 168 168 252 252
  Minus4w 168 168 252 252

> head(dat)
    Date.end N.lt Year Generation Site Company Open Open.str SessionDay Since.open  Date.open Date.start  N    Ltlog
1 2017-06-27   57 2017          1    A      SC    0  Minus0w          1          1 2017-06-26 2017-06-26 69 4.051785
2 2017-06-27   57 2017          1    B      SC    0  Minus0w          1          1 2017-06-26 2017-06-26 23 4.051785
3 2017-06-27   57 2017          1    C      SC    0  Minus0w          1          1 2017-06-26 2017-06-26 27 4.051785
4 2017-06-27   57 2017          1    D      SC    0  Minus0w          1          1 2017-06-26 2017-06-26 15 4.051785
5 2017-06-27   57 2017          1    A      SC    1  Minus1w          1          8 2017-06-19 2017-06-26 25 4.051785
6 2017-06-27   57 2017          1    B      SC    1  Minus1w          1          8 2017-06-19 2017-06-26 61 4.051785

> lure.allsites
# A tibble: 1,050 x 7
# Groups:   Year, Generation, Company, Open.str, SessionDay [1,050]
    Year Generation Company Open.str SessionDay data                N4
   <int>      <int> <chr>   <chr>         <int> <list>           <int>
 1  2017          1 SC      Minus0w           1 <tibble [4 x 8]>   134
 2  2017          1 SC      Minus1w           1 <tibble [4 x 8]>   165
 3  2017          1 SC      Minus2w           1 <tibble [4 x 8]>   149
 4  2017          1 SC      Minus3w           1 <tibble [4 x 8]>   135
 5  2017          1 SC      Minus4w           1 <tibble [4 x 8]>    99
 6  2017          1 OT      Minus0w           1 <tibble [4 x 8]>    55
 7  2017          1 OT      Minus1w           1 <tibble [4 x 8]>    56
 8  2017          1 OT      Minus2w           1 <tibble [4 x 8]>    59
 9  2017          1 OT      Minus3w           1 <tibble [4 x 8]>    25
10  2017          1 OT      Minus4w           1 <tibble [4 x 8]>    28
# ... with 1,040 more rows

> lure.lt
# A tibble: 70 x 6
# Groups:   Year, Generation, SessionDay [70]
    Year Generation SessionDay data                N.lt Company
   <int>      <int>      <int> <list>             <dbl> <chr>
 1  2017          1          1 <tibble [60 x 10]>    57 LT
 2  2017          1          2 <tibble [60 x 10]>    45 LT
 3  2017          1          3 <tibble [60 x 10]>     9 LT
 4  2017          1          4 <tibble [60 x 10]>     0 LT
 5  2017          1          5 <tibble [60 x 10]>     0 LT
 6  2017          1          6 <tibble [60 x 10]>     1 LT
 7  2017          1          7 <tibble [60 x 10]>     9 LT
 8  2017          1          8 <tibble [60 x 10]>    26 LT
 9  2017          1          9 <tibble [60 x 10]>    20 LT
10  2017          1         10 <tibble [60 x 10]>    80 LT
# ... with 60 more rows


################################################################################################################################

# Figure 1: map

#### code not shown ####

# Figure 2
# Plotting raw data


library(extrafont)
library(ggplot2)
library(ggthemes)
library(patchwork)
library(devEMF)

scale_company <- c( "OT"=hsv(343/360, 0.95, 0.82), "SC"=hsv(145/360, 0.95, 0.72), "SE"=hsv(208/360, 0.85, 0.65),
                    "LT"=hsv(0/360, 0, 0.1),
                    "Minus0w"=hsv(0.1, 0.3, 0.95), "Minus1w"=hsv(0.3, 0.3, 0.95),
                    "Minus2w"=hsv(0.5, 0.3, 0.95), "Minus3w"=hsv(0.7, 0.3, 0.95),
                    "Minus4w"=hsv(0.9, 0.3, 0.95) )
names_cpny <- c(`OT`="Lure: OT",
                `SC`="Lure: SC",
                `SE`="Lure: SE",
                `LT`="Light trap")
names_gen <- c( `1`="1st gen.",
                `2`="2nd gen.",
                `3`="3rd gen.",
                `0`="0th gen.",
                `2017`="2017",
                `2018`="2018" )

YLIM <- c(0.4, 2000)
#YLIM <- sqrt(c(0, 1700))
p3 <- ggplot(data=lure.allsites)
p3 <- p3 +
    geom_hline(yintercept=seq(1, 10, by=1), size=0.1, color=gray(0.85)) +
    geom_hline(yintercept=seq(10, 100, by=10), size=0.1, color=gray(0.85)) +
    geom_hline(yintercept=seq(100, 1000, by=100), size=0.1, color=gray(0.85)) +
    geom_hline(yintercept=c(0, 1, 10, 100, 1000), size=0.3, color=gray(0.85)) +
    geom_vline(xintercept=seq(0, 49, by=7), size=0.25, color=gray(0.85)) +
    geom_line(  aes(y=N4+0.5, x=SessionDay,
                    group=interaction(Generation, Company, Open.str), color=Company, linetype=Open.str), size=0.2  ) +
    geom_point(  aes(y=N4+0.5, x=SessionDay,
                    group=interaction(Generation, Company, Open.str), color=Company, shape=Open.str), size=0.7, stroke=0.3  ) +
    geom_line(  data=lure.lt,
                aes(y=N.lt+0.5, x=SessionDay, group=interaction(Generation), color=Company), size=0.3  ) +
    facet_grid( Company ~ Year + Generation, margins=F,
                labeller=labeller(Generation = names_gen, Company = names_cpny) ) +
    theme_classic(base_size=10) +
    scale_color_manual(name="Company", values=scale_company, guide=guide_legend(byrow=TRUE, nrow=2)) +
    scale_shape_manual(name="Open timing", values=1:5, guide=guide_legend(byrow=TRUE, nrow=2)) +
    scale_linetype_manual(name="Open timing", values=1:5, guide=guide_legend(byrow=TRUE, nrow=2)) +
    theme(  panel.background=element_blank(),
            plot.background=element_rect(color=NA, fill=NA),
            plot.margin=unit(c(0, 0, 0, 0), "lines")  ) + # top, right, bottom, left
    theme(  text=element_text(family="Arial"),
            plot.title=element_blank(),
#            plot.title=element_text(family="Arial", face="bold", size=10, angle=0, hjust=0),
            strip.background=element_blank(),
            strip.text=element_text(family="Arial", face="bold", margin=margin(b=2, t=0, l=4, r=4, unit="pt")),#, size=10
            axis.text.x=element_text(family="Arial Narrow", face="bold", size=8, angle=0, hjust=0.5),#
            axis.text.y=element_text(family="Arial Narrow", face="bold", size=8, angle=0, hjust=1)  ) + # hjust=0.5: center
    theme(  legend.position="top", legend.spacing=unit(0, "lines"),
            legend.margin=margin(b=0, t=0, l=4, r=5, unit="pt"), # the margin around each legend
            legend.box="horizontal", # align multiple legend series horizontally
            legend.box.margin=margin(b=2, t=1, l=1, r=2, unit="pt"), # margins around the full legend area
            legend.box.background=element_blank(), # do not show box around the legend
            legend.box.spacing=unit(4, unit="pt"), # space between legend box and the plot body
            legend.key.size=unit(15, "pt"), legend.key.height=unit(4, "pt"), # enlarge key but not increase margin
            legend.title=element_text(family="Arial", size=7),
            legend.text=element_text(family="Arial", face="bold", size=7)  ) +
    scale_x_continuous(expand=c(0, 1), limits=c(0, 21), breaks=seq(0, 21, by=7), labels=seq(0, 21, by=7) ) +
    scale_y_log10(expand=c(0, 0), limits=YLIM, breaks=c(1, 10, 100, 1000), labels=c(1, 10, 100, 1000)) +
    labs( title="A", x="Session day (trap settlement = 0)", y="Captured males per day + 0.5" )
ggsave("Figure2.pdf", device=cairo_pdf, plot=p3, dpi=300, width=11/2.54, height=12/2.54)
ggsave("Figure2.eps", device=cairo_pdf, plot=p3, dpi=300, width=11/2.54, height=12/2.54)
ggsave("Figure2.png", type="cairo", plot=p3, dpi=300, width=11/2.54, height=12/2.54)

emf(file="Figure2.emf", width=11/2.54, height=12/2.54)
print(p3)
dev.off()


################################################################################################################################


# Part I: Cross validation

# The mgcv package is a tool of R that fits the generalized additive model (GAM).
# Here, the function mgcv::gam() is used to fit GAMM.
# The package also has the gamm() function that fits the GAMM parameter, but random intercepts may also
# be implemented via gam() function with s(Variable, bs=“re”), where bs is the spline basis.
# The package developer also recommends gam() for simple random intercepts (Wood, 2020).


########

I (nonlinear population fluctuation, generation-specific intercept and shape)
    Generation + s(SessionDay, by=Generation)
II (nonlinear population fluctuation, generation-specific intercept)
    Generation + s(SessionDay)
III (nonlinear population fluctuation, generation-specific shape but common intercept)
    s(SessionDay, by=Generation)
IV (nonlinear population fluctuation, common intercept and shape)
    s(SessionDay)

i (population fluctuation, generation-specific intercept and slope)
    Generation * SessionDay
ii (population fluctuation, generation-specific intercept)
    Generation + SessionDay
iii (population fluctuation, generation-specific slope but common slope)
    Generation:SessionDay
iv (population fluctuation, common intercept and shape)
    SessionDay

I` (nonlinear population fluctuation approximated by light trap, generation-specific intercept and shape)
    Generation + s(Ltlog, by=Generation)
II` (nonlinear population fluctuation approximated by light trap, generation-specific intercept)
    Generation + s(Ltlog)
III` (nonlinear population fluctuation approximated by light trap, generation-specific shape but common intercept)
    s(Ltlog, by=Generation)
IV` (nonlinear population fluctuation approximated by light trap, common intercept and shape)
    s(Ltlog)

i` (population fluctuation approximated by light trap, generation-specific intercept and slope)
    Generation * Ltlog
ii` (population fluctuation approximated by light trap, generation-specific intercept)
    Generation + Ltlog
iii` (population fluctuation approximated by light trap, generation-specific slope but common slope)
    Generation:Ltlog
iv` (population fluctuation approximated by light trap, common intercept and shape)
    Ltlog

P (nonlinear attenuation, company-specific intercept and shape)
    Company + s(Since.open, by=Company)
Q (linear attenuation, company-specific intercept and slope)
    Company * Since.open

########


# first, make table.
is.LT <- rep(c(FALSE, TRUE), each=16)
is.dyn.lin <- c(rep(FALSE, 4), rep(TRUE, 4)) %>% rep(each=2) %>% rep(2)
is.com.itcpt <- c(rep(FALSE, 4), rep(TRUE, 4)) %>% rep(2) %>% rep(2)
is.com.slope <- c(FALSE, TRUE) %>% rep(each=2) %>% rep(2) %>% rep(2) %>% rep(2)
is.att.lin <- rep(c(FALSE, TRUE), 16)
gam.sday.formula <- c(  "Generation + s(SessionDay, by=Generation)",
                        "Generation + s(SessionDay)",
                        "s(SessionDay, by=Generation)",
                        "s(SessionDay)"  )
lin.sday.formula <- c(  "Generation * SessionDay",
                        "Generation + SessionDay",
                        "Generation:SessionDay",
                        "SessionDay"  )
gam.lt.formula <- c(    "Generation + s(Ltlog, by=Generation)",
                        "Generation + s(Ltlog)",
                        "s(Ltlog, by=Generation)",
                        "s(Ltlog)"  )
lin.lt.formula <- c(    "Generation * Ltlog",
                        "Generation + Ltlog",
                        "Generation:Ltlog",
                        "Ltlog"  )
since.formula <- c( "Company + s(Since.open, by=Company)",
                    "Company *  Since.open" )

tib.total <- tidyr::expand_grid(frm.lhs="N ~",
                                frm.dyn=c(gam.sday.formula, lin.sday.formula, gam.lt.formula, lin.lt.formula),
                                frm.pls="+",
                                frm.att=since.formula) %>%
    dplyr::mutate(  frm.str=paste(frm.lhs, frm.dyn, frm.pls, frm.att, sep=" ") ) %>%
    dplyr::mutate(  is.LT=is.LT,
                    is.dyn.lin=is.dyn.lin,
                    is.com.itcpt=is.com.itcpt,
                    is.com.slope=is.com.slope,
                    is.att.lin=is.att.lin  ) %>%
    dplyr::mutate(  result=list(0)  ) %>%
    dplyr::mutate(  num=1:nrow(.)  )
tib.total


# just run to check the labels
tib.total %>% dplyr::select(num, starts_with("is"), "frm.dyn") %>% data.frame()

# just run to check the labels for base dynamics
tib.total %>%
    dplyr::select(num, starts_with("is"), "frm.dyn") %>%
    dplyr::arrange(is.com.slope, is.com.itcpt, is.dyn.lin, is.LT) %>%
    data.frame()

# just run to check the labels for lure attenuation
tib.total %>%
    dplyr::select(num, starts_with("is"), "frm.att") %>%
    dplyr::arrange(is.att.lin) %>%
    data.frame()


> tib.total
# A tibble: 32 x 12
   frm.lhs frm.dyn     frm.pls frm.att   frm.str        is.LT is.com.itcpt is.com.slope is.dyn.lin is.att.lin result   num
   <chr>   <chr>       <chr>   <chr>     <chr>          <lgl> <lgl>        <lgl>        <lgl>      <lgl>      <list> <int>
 1 N ~     Generation~ +       Company ~ N ~ Generatio~ FALSE FALSE        FALSE        FALSE      FALSE      <dbl ~     1
 2 N ~     Generation~ +       Company ~ N ~ Generatio~ FALSE FALSE        FALSE        FALSE      TRUE       <dbl ~     2
 3 N ~     Generation~ +       Company ~ N ~ Generatio~ FALSE FALSE        TRUE         FALSE      FALSE      <dbl ~     3
 4 N ~     Generation~ +       Company ~ N ~ Generatio~ FALSE FALSE        TRUE         FALSE      TRUE       <dbl ~     4
 5 N ~     s(SessionD~ +       Company ~ N ~ s(Session~ FALSE TRUE         FALSE        FALSE      FALSE      <dbl ~     5
 6 N ~     s(SessionD~ +       Company ~ N ~ s(Session~ FALSE TRUE         FALSE        FALSE      TRUE       <dbl ~     6
 7 N ~     s(SessionD~ +       Company ~ N ~ s(Session~ FALSE TRUE         TRUE         FALSE      FALSE      <dbl ~     7
 8 N ~     s(SessionD~ +       Company ~ N ~ s(Session~ FALSE TRUE         TRUE         FALSE      TRUE       <dbl ~     8
 9 N ~     Generation~ +       Company ~ N ~ Generatio~ FALSE FALSE        FALSE        TRUE       FALSE      <dbl ~     9
10 N ~     Generation~ +       Company ~ N ~ Generatio~ FALSE FALSE        FALSE        TRUE       TRUE       <dbl ~    10
# ... with 22 more rows

> tib.total %>% dplyr::select(num, starts_with("is"), "frm.dyn") %>% data.frame()
   num is.LT is.dyn.lin is.com.itcpt is.com.slope is.att.lin                                   frm.dyn
1    1 FALSE      FALSE        FALSE        FALSE      FALSE Generation + s(SessionDay, by=Generation)
2    2 FALSE      FALSE        FALSE        FALSE       TRUE Generation + s(SessionDay, by=Generation)
3    3 FALSE      FALSE        FALSE         TRUE      FALSE                Generation + s(SessionDay)
4    4 FALSE      FALSE        FALSE         TRUE       TRUE                Generation + s(SessionDay)
5    5 FALSE      FALSE         TRUE        FALSE      FALSE              s(SessionDay, by=Generation)
6    6 FALSE      FALSE         TRUE        FALSE       TRUE              s(SessionDay, by=Generation)
7    7 FALSE      FALSE         TRUE         TRUE      FALSE                             s(SessionDay)
8    8 FALSE      FALSE         TRUE         TRUE       TRUE                             s(SessionDay)
9    9 FALSE       TRUE        FALSE        FALSE      FALSE                   Generation * SessionDay
10  10 FALSE       TRUE        FALSE        FALSE       TRUE                   Generation * SessionDay
11  11 FALSE       TRUE        FALSE         TRUE      FALSE                   Generation + SessionDay
12  12 FALSE       TRUE        FALSE         TRUE       TRUE                   Generation + SessionDay
13  13 FALSE       TRUE         TRUE        FALSE      FALSE                     Generation:SessionDay
14  14 FALSE       TRUE         TRUE        FALSE       TRUE                     Generation:SessionDay
15  15 FALSE       TRUE         TRUE         TRUE      FALSE                                SessionDay
16  16 FALSE       TRUE         TRUE         TRUE       TRUE                                SessionDay
17  17  TRUE      FALSE        FALSE        FALSE      FALSE      Generation + s(Ltlog, by=Generation)
18  18  TRUE      FALSE        FALSE        FALSE       TRUE      Generation + s(Ltlog, by=Generation)
19  19  TRUE      FALSE        FALSE         TRUE      FALSE                     Generation + s(Ltlog)
20  20  TRUE      FALSE        FALSE         TRUE       TRUE                     Generation + s(Ltlog)
21  21  TRUE      FALSE         TRUE        FALSE      FALSE                   s(Ltlog, by=Generation)
22  22  TRUE      FALSE         TRUE        FALSE       TRUE                   s(Ltlog, by=Generation)
23  23  TRUE      FALSE         TRUE         TRUE      FALSE                                  s(Ltlog)
24  24  TRUE      FALSE         TRUE         TRUE       TRUE                                  s(Ltlog)
25  25  TRUE       TRUE        FALSE        FALSE      FALSE                        Generation * Ltlog
26  26  TRUE       TRUE        FALSE        FALSE       TRUE                        Generation * Ltlog
27  27  TRUE       TRUE        FALSE         TRUE      FALSE                        Generation + Ltlog
28  28  TRUE       TRUE        FALSE         TRUE       TRUE                        Generation + Ltlog
29  29  TRUE       TRUE         TRUE        FALSE      FALSE                          Generation:Ltlog
30  30  TRUE       TRUE         TRUE        FALSE       TRUE                          Generation:Ltlog
31  31  TRUE       TRUE         TRUE         TRUE      FALSE                                     Ltlog
32  32  TRUE       TRUE         TRUE         TRUE       TRUE                                     Ltlog

> tib.total %>%
+     dplyr::select(num, starts_with("is"), "frm.dyn") %>%
+     dplyr::arrange(is.com.slope, is.com.itcpt, is.dyn.lin, is.LT) %>%
+     data.frame()
   num is.LT is.dyn.lin is.com.itcpt is.com.slope is.att.lin                                   frm.dyn
1    1 FALSE      FALSE        FALSE        FALSE      FALSE Generation + s(SessionDay, by=Generation)
2    2 FALSE      FALSE        FALSE        FALSE       TRUE Generation + s(SessionDay, by=Generation)
3   17  TRUE      FALSE        FALSE        FALSE      FALSE      Generation + s(Ltlog, by=Generation)
4   18  TRUE      FALSE        FALSE        FALSE       TRUE      Generation + s(Ltlog, by=Generation)
5    9 FALSE       TRUE        FALSE        FALSE      FALSE                   Generation * SessionDay
6   10 FALSE       TRUE        FALSE        FALSE       TRUE                   Generation * SessionDay
7   25  TRUE       TRUE        FALSE        FALSE      FALSE                        Generation * Ltlog
8   26  TRUE       TRUE        FALSE        FALSE       TRUE                        Generation * Ltlog
9    5 FALSE      FALSE         TRUE        FALSE      FALSE              s(SessionDay, by=Generation)
10   6 FALSE      FALSE         TRUE        FALSE       TRUE              s(SessionDay, by=Generation)
11  21  TRUE      FALSE         TRUE        FALSE      FALSE                   s(Ltlog, by=Generation)
12  22  TRUE      FALSE         TRUE        FALSE       TRUE                   s(Ltlog, by=Generation)
13  13 FALSE       TRUE         TRUE        FALSE      FALSE                     Generation:SessionDay
14  14 FALSE       TRUE         TRUE        FALSE       TRUE                     Generation:SessionDay
15  29  TRUE       TRUE         TRUE        FALSE      FALSE                          Generation:Ltlog
16  30  TRUE       TRUE         TRUE        FALSE       TRUE                          Generation:Ltlog
17   3 FALSE      FALSE        FALSE         TRUE      FALSE                Generation + s(SessionDay)
18   4 FALSE      FALSE        FALSE         TRUE       TRUE                Generation + s(SessionDay)
19  19  TRUE      FALSE        FALSE         TRUE      FALSE                     Generation + s(Ltlog)
20  20  TRUE      FALSE        FALSE         TRUE       TRUE                     Generation + s(Ltlog)
21  11 FALSE       TRUE        FALSE         TRUE      FALSE                   Generation + SessionDay
22  12 FALSE       TRUE        FALSE         TRUE       TRUE                   Generation + SessionDay
23  27  TRUE       TRUE        FALSE         TRUE      FALSE                        Generation + Ltlog
24  28  TRUE       TRUE        FALSE         TRUE       TRUE                        Generation + Ltlog
25   7 FALSE      FALSE         TRUE         TRUE      FALSE                             s(SessionDay)
26   8 FALSE      FALSE         TRUE         TRUE       TRUE                             s(SessionDay)
27  23  TRUE      FALSE         TRUE         TRUE      FALSE                                  s(Ltlog)
28  24  TRUE      FALSE         TRUE         TRUE       TRUE                                  s(Ltlog)
29  15 FALSE       TRUE         TRUE         TRUE      FALSE                                SessionDay
30  16 FALSE       TRUE         TRUE         TRUE       TRUE                                SessionDay
31  31  TRUE       TRUE         TRUE         TRUE      FALSE                                     Ltlog
32  32  TRUE       TRUE         TRUE         TRUE       TRUE                                     Ltlog

> # just run to check the labels for lure attenuation
> tib.total %>%
+     dplyr::select(num, starts_with("is"), "frm.att") %>%
+     dplyr::arrange(is.att.lin) %>%
+     data.frame()
   num is.LT is.dyn.lin is.com.itcpt is.com.slope is.att.lin                             frm.att
1    1 FALSE      FALSE        FALSE        FALSE      FALSE Company + s(Since.open, by=Company)
2    3 FALSE      FALSE        FALSE         TRUE      FALSE Company + s(Since.open, by=Company)
3    5 FALSE      FALSE         TRUE        FALSE      FALSE Company + s(Since.open, by=Company)
4    7 FALSE      FALSE         TRUE         TRUE      FALSE Company + s(Since.open, by=Company)
5    9 FALSE       TRUE        FALSE        FALSE      FALSE Company + s(Since.open, by=Company)
6   11 FALSE       TRUE        FALSE         TRUE      FALSE Company + s(Since.open, by=Company)
7   13 FALSE       TRUE         TRUE        FALSE      FALSE Company + s(Since.open, by=Company)
8   15 FALSE       TRUE         TRUE         TRUE      FALSE Company + s(Since.open, by=Company)
9   17  TRUE      FALSE        FALSE        FALSE      FALSE Company + s(Since.open, by=Company)
10  19  TRUE      FALSE        FALSE         TRUE      FALSE Company + s(Since.open, by=Company)
11  21  TRUE      FALSE         TRUE        FALSE      FALSE Company + s(Since.open, by=Company)
12  23  TRUE      FALSE         TRUE         TRUE      FALSE Company + s(Since.open, by=Company)
13  25  TRUE       TRUE        FALSE        FALSE      FALSE Company + s(Since.open, by=Company)
14  27  TRUE       TRUE        FALSE         TRUE      FALSE Company + s(Since.open, by=Company)
15  29  TRUE       TRUE         TRUE        FALSE      FALSE Company + s(Since.open, by=Company)
16  31  TRUE       TRUE         TRUE         TRUE      FALSE Company + s(Since.open, by=Company)
17   2 FALSE      FALSE        FALSE        FALSE       TRUE               Company *  Since.open
18   4 FALSE      FALSE        FALSE         TRUE       TRUE               Company *  Since.open
19   6 FALSE      FALSE         TRUE        FALSE       TRUE               Company *  Since.open
20   8 FALSE      FALSE         TRUE         TRUE       TRUE               Company *  Since.open
21  10 FALSE       TRUE        FALSE        FALSE       TRUE               Company *  Since.open
22  12 FALSE       TRUE        FALSE         TRUE       TRUE               Company *  Since.open
23  14 FALSE       TRUE         TRUE        FALSE       TRUE               Company *  Since.open
24  16 FALSE       TRUE         TRUE         TRUE       TRUE               Company *  Since.open
25  18  TRUE      FALSE        FALSE        FALSE       TRUE               Company *  Since.open
26  20  TRUE      FALSE        FALSE         TRUE       TRUE               Company *  Since.open
27  22  TRUE      FALSE         TRUE        FALSE       TRUE               Company *  Since.open
28  24  TRUE      FALSE         TRUE         TRUE       TRUE               Company *  Since.open
29  26  TRUE       TRUE        FALSE        FALSE       TRUE               Company *  Since.open
30  28  TRUE       TRUE        FALSE         TRUE       TRUE               Company *  Since.open
31  30  TRUE       TRUE         TRUE        FALSE       TRUE               Company *  Since.open
32  32  TRUE       TRUE         TRUE         TRUE       TRUE               Company *  Since.open

################################################################################################################################

# fitting the candidate models. It takes several minutes.

# assuming negative binomial distribution to deal with overdispersion
# https://stat.ethz.ch/R-manual/R-patched/library/mgcv/html/negbin.html


for (i in 1:nrow(tib.total)) {
    frm.str <- purrr::pluck(tib.total, "frm.str", i)
    print(frm.str)
    ptime0 <- proc.time()
    try({
        tib.total[i, "result"][[1]] <- dat %>%
            tidyr::expand_grid(kf.drop=as.factor(c("A", "B", "C", "D"))) %>%
            dplyr::group_by(kf.drop) %>%
            tidyr::nest() %>%
            dplyr::mutate(  data.test=purrr::map(.x=data, .f=~dplyr::filter(., Site==kf.drop))) %>%
            dplyr::mutate(  data.train=purrr::map(.x=data, .f=~dplyr::filter(., Site!=kf.drop))) %>%
            dplyr::mutate(  gam=purrr::map(.x=data.train, .f=~mgcv::gam(formula=as.formula(frm.str),
                                                                        data=., family=nb(theta=NULL, link="log") ))  ) %>%
            dplyr::mutate(  aic=purrr::map_dbl(.x=gam, .f=~AIC(.))  ) %>%
            dplyr::mutate(  gcv=purrr::map_dbl(.x=gam, .f=~pluck(., "gcv.ubre"))  ) %>%
            dplyr::mutate(  dev.expl=purrr::map_dbl(  .x=gam, .f=~pluck(summary(.), "dev.expl"))  ) %>%
            dplyr::mutate(  pred=purrr::map(  .x=gam, .f=~predict(object=., newdata=pluck(data.test, 1), type="response"))  ) %>%
            dplyr::mutate(  RMSE=sqrt(sum((pluck(data.test, 1, "N")-pluck(pred, 1))^2)/length(pluck(pred, 1)))  ) %>%
            dplyr::mutate(  NLL=-sum(dpois(  pluck(data.test, 1, "N"), pluck(pred, 1), log=TRUE  ))  ) %>%
            list()
    })
    cal.time <- proc.time()-ptime0
    print(cal.time)
}

# see the whole results
tib.total$result

# calculate sums of NLL
tib.total <- tib.total %>%
    dplyr::mutate(sum.NLL=purrr::map_dbl(.x=result, .f=~(purrr::pluck(., "NLL") %>% sum(.))))

# print the NLL sums
tib.arrange <- tib.total %>%
    dplyr::arrange(sum.NLL) %>%
    dplyr::mutate(delta.NLL=sum.NLL - min(sum.NLL)) %>%
    dplyr::select(sum.NLL, delta.NLL, num, frm.str) %>%
    data.frame()
tib.arrange

# print the NLL sums (by conditions)
tib.arrange2 <- tib.total %>%
    dplyr::mutate(delta.NLL=sum.NLL - min(sum.NLL)) %>%
    dplyr::select(num, starts_with("is"), sum.NLL, delta.NLL) %>%
    dplyr::arrange(is.com.itcpt, is.com.slope, is.LT, is.dyn.lin, is.att.lin) %>%
    data.frame()
tib.arrange2

################################################################################################################################

# smaller negative log likelihood (NLL) = good generalization performance for unknown test site.

> tib.arrange
    sum.NLL  delta.NLL num                                                                             frm.str
1  32555.05     0.0000   2               N ~ Generation + s(SessionDay, by=Generation) + Company *  Since.open
2  32669.37   114.3232   1 N ~ Generation + s(SessionDay, by=Generation) + Company + s(Since.open, by=Company)
3  33909.49  1354.4412   6                            N ~ s(SessionDay, by=Generation) + Company *  Since.open
4  34057.05  1501.9979   5              N ~ s(SessionDay, by=Generation) + Company + s(Since.open, by=Company)
5  36997.72  4442.6716  18                    N ~ Generation + s(Ltlog, by=Generation) + Company *  Since.open
6  37102.57  4547.5212  22                                 N ~ s(Ltlog, by=Generation) + Company *  Since.open
7  37142.21  4587.1643  17      N ~ Generation + s(Ltlog, by=Generation) + Company + s(Since.open, by=Company)
8  37326.94  4771.8934  21                   N ~ s(Ltlog, by=Generation) + Company + s(Since.open, by=Company)
9  40345.06  7790.0091   9                   N ~ Generation * SessionDay + Company + s(Since.open, by=Company)
10 40424.87  7869.8172  10                                 N ~ Generation * SessionDay + Company *  Since.open
11 40643.16  8088.1108  26                                      N ~ Generation * Ltlog + Company *  Since.open
12 40685.74  8130.6896  20                                   N ~ Generation + s(Ltlog) + Company *  Since.open
13 40875.95  8320.9050  25                        N ~ Generation * Ltlog + Company + s(Since.open, by=Company)
14 40926.33  8371.2815  19                     N ~ Generation + s(Ltlog) + Company + s(Since.open, by=Company)
15 42360.42  9805.3715  28                                      N ~ Generation + Ltlog + Company *  Since.open
16 42516.23  9961.1763  27                        N ~ Generation + Ltlog + Company + s(Since.open, by=Company)
17 42622.42 10067.3673   4                              N ~ Generation + s(SessionDay) + Company *  Since.open
18 42709.21 10154.1625   3                N ~ Generation + s(SessionDay) + Company + s(Since.open, by=Company)
19 44814.04 12258.9911  30                                        N ~ Generation:Ltlog + Company *  Since.open
20 44861.90 12306.8532  29                          N ~ Generation:Ltlog + Company + s(Since.open, by=Company)
21 44925.89 12370.8393   8                                           N ~ s(SessionDay) + Company *  Since.open
22 45048.07 12493.0227   7                             N ~ s(SessionDay) + Company + s(Since.open, by=Company)
23 45304.30 12749.2535  13                     N ~ Generation:SessionDay + Company + s(Since.open, by=Company)
24 45337.20 12782.1558  14                                   N ~ Generation:SessionDay + Company *  Since.open
25 45575.07 13020.0247  24                                                N ~ s(Ltlog) + Company *  Since.open
26 45732.37 13177.3176  23                                  N ~ s(Ltlog) + Company + s(Since.open, by=Company)
27 45757.72 13202.6683  12                                 N ~ Generation + SessionDay + Company *  Since.open
28 45773.11 13218.0646  11                   N ~ Generation + SessionDay + Company + s(Since.open, by=Company)
29 48000.14 15445.0953  16                                              N ~ SessionDay + Company *  Since.open
30 48071.87 15516.8192  15                                N ~ SessionDay + Company + s(Since.open, by=Company)
31 48766.96 16211.9127  32                                                   N ~ Ltlog + Company *  Since.open
32 48854.79 16299.7453  31                                     N ~ Ltlog + Company + s(Since.open, by=Company)

> tib.arrange2
   num is.LT is.dyn.lin is.com.itcpt is.com.slope is.att.lin  sum.NLL  delta.NLL
1    1 FALSE      FALSE        FALSE        FALSE      FALSE 32669.37   114.3232
2    2 FALSE      FALSE        FALSE        FALSE       TRUE 32555.05     0.0000
3    9 FALSE       TRUE        FALSE        FALSE      FALSE 40345.06  7790.0091
4   10 FALSE       TRUE        FALSE        FALSE       TRUE 40424.87  7869.8172
5   17  TRUE      FALSE        FALSE        FALSE      FALSE 37142.21  4587.1643
6   18  TRUE      FALSE        FALSE        FALSE       TRUE 36997.72  4442.6716
7   25  TRUE       TRUE        FALSE        FALSE      FALSE 40875.95  8320.9050
8   26  TRUE       TRUE        FALSE        FALSE       TRUE 40643.16  8088.1108
9    3 FALSE      FALSE        FALSE         TRUE      FALSE 42709.21 10154.1625
10   4 FALSE      FALSE        FALSE         TRUE       TRUE 42622.42 10067.3673
11  11 FALSE       TRUE        FALSE         TRUE      FALSE 45773.11 13218.0646
12  12 FALSE       TRUE        FALSE         TRUE       TRUE 45757.72 13202.6683
13  19  TRUE      FALSE        FALSE         TRUE      FALSE 40926.33  8371.2815
14  20  TRUE      FALSE        FALSE         TRUE       TRUE 40685.74  8130.6896
15  27  TRUE       TRUE        FALSE         TRUE      FALSE 42516.23  9961.1763
16  28  TRUE       TRUE        FALSE         TRUE       TRUE 42360.42  9805.3715
17   5 FALSE      FALSE         TRUE        FALSE      FALSE 34057.05  1501.9979
18   6 FALSE      FALSE         TRUE        FALSE       TRUE 33909.49  1354.4412
19  13 FALSE       TRUE         TRUE        FALSE      FALSE 45304.30 12749.2535
20  14 FALSE       TRUE         TRUE        FALSE       TRUE 45337.20 12782.1558
21  21  TRUE      FALSE         TRUE        FALSE      FALSE 37326.94  4771.8934
22  22  TRUE      FALSE         TRUE        FALSE       TRUE 37102.57  4547.5212
23  29  TRUE       TRUE         TRUE        FALSE      FALSE 44861.90 12306.8532
24  30  TRUE       TRUE         TRUE        FALSE       TRUE 44814.04 12258.9911
25   7 FALSE      FALSE         TRUE         TRUE      FALSE 45048.07 12493.0227
26   8 FALSE      FALSE         TRUE         TRUE       TRUE 44925.89 12370.8393
27  15 FALSE       TRUE         TRUE         TRUE      FALSE 48071.87 15516.8192
28  16 FALSE       TRUE         TRUE         TRUE       TRUE 48000.14 15445.0953
29  23  TRUE      FALSE         TRUE         TRUE      FALSE 45732.37 13177.3176
30  24  TRUE      FALSE         TRUE         TRUE       TRUE 45575.07 13020.0247
31  31  TRUE       TRUE         TRUE         TRUE      FALSE 48854.79 16299.7453
32  32  TRUE       TRUE         TRUE         TRUE       TRUE 48766.96 16211.9127

################################################################################################################################

> tib.total$result
[[1]]
# A tibble: 4 x 11
# Groups:   kf.drop [4]
  kf.drop data                data.test           data.train           gam      aic   gcv dev.expl pred         RMSE   NLL
  <fct>   <list>              <list>              <list>               <lis>  <dbl> <dbl>    <dbl> <list>      <dbl> <dbl>
1 A       <tibble [4,200 x 1~ <tibble [1,050 x 1~ <tibble [3,150 x 14~ <gam> 18852. 9490.    0.535 <dbl [1,05~  23.6 8786.
2 B       <tibble [4,200 x 1~ <tibble [1,050 x 1~ <tibble [3,150 x 14~ <gam> 19395. 9786.    0.567 <dbl [1,05~  17.3 6579.
3 C       <tibble [4,200 x 1~ <tibble [1,050 x 1~ <tibble [3,150 x 14~ <gam> 18292. 9232.    0.583 <dbl [1,05~  22.3 9812.
4 D       <tibble [4,200 x 1~ <tibble [1,050 x 1~ <tibble [3,150 x 14~ <gam> 19614. 9893.    0.603 <dbl [1,05~  18.7 7493.

[[2]]
# A tibble: 4 x 11
# Groups:   kf.drop [4]
  kf.drop data                data.test           data.train           gam      aic   gcv dev.expl pred         RMSE   NLL
  <fct>   <list>              <list>              <list>               <lis>  <dbl> <dbl>    <dbl> <list>      <dbl> <dbl>
1 A       <tibble [4,200 x 1~ <tibble [1,050 x 1~ <tibble [3,150 x 14~ <gam> 18891. 9513.    0.527 <dbl [1,05~  23.7 8768.
2 B       <tibble [4,200 x 1~ <tibble [1,050 x 1~ <tibble [3,150 x 14~ <gam> 19457. 9821.    0.556 <dbl [1,05~  16.9 6363.
3 C       <tibble [4,200 x 1~ <tibble [1,050 x 1~ <tibble [3,150 x 14~ <gam> 18325. 9254.    0.577 <dbl [1,05~  22.7 9813.
4 D       <tibble [4,200 x 1~ <tibble [1,050 x 1~ <tibble [3,150 x 14~ <gam> 19646. 9917.    0.597 <dbl [1,05~  19.3 7610.

... 32 models in all


################################################################################################################################

################################################################################################################################


# Part II: model with pheromone lures
# the final model with mgcv::gam()

#### M2: linear attenuation in the attractiveness after the lure exposure to air.
#### single smoother term for male density fluctuation
ptime0 <- proc.time()
nest.gamm.M2 <- dat %>%
    tidyr::nest(data=everything()) %>%
    dplyr::mutate(  gam=purrr::map(.x=data, .f=~mgcv::gam(  N ~ s(Site, bs="re") +
                                                            Generation + s(SessionDay, by=Generation, bs="tp") +
                                                            Company*Since.open,
                                                            data=., family=nb(theta=NULL, link="log") ))  ) %>%
    dplyr::mutate(  aic=purrr::map_dbl(.x=gam, .f=~AIC(.))  ) %>%
    dplyr::mutate(  gcv=purrr::map_dbl(.x=gam, .f=~pluck(., "gcv.ubre"))  ) %>%
    dplyr::mutate(  coef=purrr::map(.x=gam, .f=~coef(.))  ) %>%
    dplyr::mutate(  intercept=purrr::map_dbl(.x=coef, .f=~pluck(., "(Intercept)"))  ) %>%
    dplyr::mutate(  deviance=purrr::map(  .x=gam, .f=~residuals(., type="deviance"))  ) %>%
    dplyr::mutate(  resid.work=purrr::map(  .x=gam, .f=~residuals(., type="working"))  ) %>%
    dplyr::mutate(  weights.work=purrr::map(.x=gam, .f=~pluck(., "weights"))  ) %>%
    dplyr::mutate(  pred.link=purrr::map(   .x=gam,
                                            .f=~as_tibble(data.frame(predict(., type="link", se.fit=TRUE)))   )  ) %>%
    dplyr::mutate(  pred.terms=purrr::map(  .x=gam,
                                            .f=~as_tibble(data.frame(predict(., type="terms", se.fit=TRUE)))  )  )
cal.time <- proc.time()-ptime0; print(cal.time);
nest.gamm.M2
purrr::map(pluck(nest.gamm.M2, "gam"), ~summary(.))


colnames(nest.gamm.M2)
unnest.gamm.M2 <- nest.gamm.M2 %>%
    dplyr::select(-gam, -aic, -gcv, -coef) %>%
    tidyr::unnest(c(data, intercept, deviance, resid.work, weights.work, pred.link, pred.terms)) %>%
    dplyr::ungroup() %>%
    dplyr::mutate(  fit.s.SessionDay..Generation =  fit.s.SessionDay..Generation0+fit.s.SessionDay..Generation1+
                                                    fit.s.SessionDay..Generation2+fit.s.SessionDay..Generation3,
                    se.fit.s.SessionDay..Generation=se.fit.s.SessionDay..Generation0+se.fit.s.SessionDay..Generation1+
                                                    se.fit.s.SessionDay..Generation2+se.fit.s.SessionDay..Generation3  )
unnest.gamm.M2
colnames(unnest.gamm.M2)


#### M1: non-linear attenuation in the attractiveness after the lure exposure to air.
#### two smoother terms for male density fluctuation and lure attenuation
ptime0 <- proc.time()
nest.gamm.M1 <- dat %>%
    tidyr::nest(data=everything()) %>%
    dplyr::mutate(  gam=purrr::map(.x=data, .f=~mgcv::gam(  N ~ s(Site, bs="re") +
                                                            Generation + s(SessionDay, by=Generation, bs="tp") +
                                                            Company + s(Since.open, by=Company, bs="tp"),
                                                            data=., family=nb(theta=NULL, link="log") ))  ) %>%
    dplyr::mutate(  aic=purrr::map_dbl(.x=gam, .f=~AIC(.))  ) %>%
    dplyr::mutate(  gcv=purrr::map_dbl(.x=gam, .f=~pluck(., "gcv.ubre"))  ) %>%
    dplyr::mutate(  coef=purrr::map(.x=gam, .f=~coef(.))  ) %>%
    dplyr::mutate(  intercept=purrr::map_dbl(.x=coef, .f=~pluck(., "(Intercept)"))  ) %>%
    dplyr::mutate(  deviance=purrr::map(  .x=gam, .f=~residuals(., type="deviance"))  ) %>%
    dplyr::mutate(  resid.work=purrr::map(  .x=gam, .f=~residuals(., type="working"))  ) %>%
    dplyr::mutate(  weights.work=purrr::map(.x=gam, .f=~pluck(., "weights"))  ) %>%
    dplyr::mutate(  pred.link=purrr::map(   .x=gam,
                                            .f=~as_tibble(data.frame(predict(., type="link", se.fit=TRUE)))   )  ) %>%
    dplyr::mutate(  pred.terms=purrr::map(  .x=gam,
                                            .f=~as_tibble(data.frame(predict(., type="terms", se.fit=TRUE)))  )  )
cal.time <- proc.time()-ptime0; print(cal.time);
nest.gamm.M1
purrr::map(pluck(nest.gamm.M1, "gam"), ~summary(.))

colnames(nest.gamm.M1)
unnest.gamm.M1 <- nest.gamm.M1 %>%
    dplyr::select(-gam, -aic, -gcv, -coef) %>%
    tidyr::unnest(c(data, intercept, deviance, resid.work, weights.work, pred.link, pred.terms)) %>%
    dplyr::ungroup() %>%
    dplyr::mutate(  fit.s.SessionDay..Generation =  fit.s.SessionDay..Generation0+fit.s.SessionDay..Generation1+
                                                    fit.s.SessionDay..Generation2+fit.s.SessionDay..Generation3,
                    se.fit.s.SessionDay..Generation=se.fit.s.SessionDay..Generation0+se.fit.s.SessionDay..Generation1+
                                                    se.fit.s.SessionDay..Generation2+se.fit.s.SessionDay..Generation3  ) %>%
    dplyr::mutate(  fit.s.Since.open..Company =     fit.s.Since.open..CompanyOT+fit.s.Since.open..CompanySC+
                                                    fit.s.Since.open..CompanySE,
                    se.fit.s.Since.open..Company =  se.fit.s.Since.open..CompanyOT+se.fit.s.Since.open..CompanySC+
                                                    se.fit.s.Since.open..CompanySE,  )
unnest.gamm.M1
colnames(unnest.gamm.M1)


# print AIC values
nest.gamm.M2$aic
nest.gamm.M1$aic

nest.gamm.M2$aic-nest.gamm.M1$aic

# plot the profiles of smoother terms
pdf("GAM_smoother_profile.pdf")
purrr::walk(.x=pluck(nest.gamm.M2, "gam"), .f=~plot(., ylim=c(-10, 10), residuals=TRUE, rug=TRUE, se=TRUE,
                                                        main="gamm.M2", pages=1, col=7))
purrr::walk(.x=pluck(nest.gamm.M1, "gam"), .f=~plot(., ylim=c(-10, 10), residuals=TRUE, rug=TRUE, se=TRUE,
                                                        main="gamm.M1", pages=2, col=6))
purrr::walk(.x=pluck(nest.gamm.M2, "gam"), .f=~gam.check(.))
purrr::walk(.x=pluck(nest.gamm.M1, "gam"), .f=~gam.check(.))
dev.off()


################################################################################################################################


> nest.gamm.M2
# A tibble: 1 x 11
 data          gam       aic    gcv coef     intercept deviance   resid.work  weights.work pred.link      pred.terms
 <list>        <list>  <dbl>  <dbl> <list>       <dbl> <list>     <list>      <list>       <list>         <list>
1 <tibble [4,2~ <gam>  25072. 12642. <dbl [4~      136. <dbl [4,2~ <dbl [4,20~ <dbl [4,200~ <tibble [4,20~ <tibble [4,200~


> purrr::map(pluck(nest.gamm.M2, "gam"), ~summary(.))
[[1]]

Family: Negative Binomial(1.434)
Link function: log

Formula:
N ~ s(Site, bs = "re") + Generation + s(SessionDay, by = Generation,
   bs = "tp") + Company * Since.open

Parametric coefficients:
                      Estimate Std. Error z value Pr(>|z|)
(Intercept)           1.358e+02  2.979e+01   4.557 5.20e-06 ***
Generation1          -1.348e+02  2.980e+01  -4.524 6.08e-06 ***
Generation2          -1.345e+02  2.979e+01  -4.515 6.32e-06 ***
Generation3          -1.337e+02  2.979e+01  -4.488 7.20e-06 ***
CompanySC             5.126e-01  8.489e-02   6.038 1.56e-09 ***
CompanySE             1.334e+00  8.303e-02  16.061  < 2e-16 ***
Since.open           -2.564e-02  2.597e-03  -9.873  < 2e-16 ***
CompanySC:Since.open  1.110e-02  3.375e-03   3.289  0.00101 **
CompanySE:Since.open  7.713e-03  3.295e-03   2.341  0.01925 *
---
Signif. codes:  0 ‘***’ 0.001 ‘**’ 0.01 ‘*’ 0.05 ‘.’ 0.1 ‘ ’ 1

Approximate significance of smooth terms:
                           edf Ref.df Chi.sq p-value
s(Site)                   2.979  3.000  423.1  <2e-16 ***
s(SessionDay):Generation0 8.094  8.233  722.1  <2e-16 ***
s(SessionDay):Generation1 6.238  6.611  756.9  <2e-16 ***
s(SessionDay):Generation2 8.474  8.921  728.7  <2e-16 ***
s(SessionDay):Generation3 8.683  8.971  816.7  <2e-16 ***
---
Signif. codes:  0 ‘***’ 0.001 ‘**’ 0.01 ‘*’ 0.05 ‘.’ 0.1 ‘ ’ 1

R-sq.(adj) =  0.326   Deviance explained = 60.2%
-REML =  12642  Scale est. = 1         n = 4200


> nest.gamm.M1
# A tibble: 1 x 11
 data          gam       aic    gcv coef     intercept deviance   resid.work  weights.work pred.link      pred.terms
 <list>        <list>  <dbl>  <dbl> <list>       <dbl> <list>     <list>      <list>       <list>         <list>
1 <tibble [4,2~ <gam>  25031. 12615. <dbl [7~      135. <dbl [4,2~ <dbl [4,20~ <dbl [4,200~ <tibble [4,20~ <tibble [4,200~


> purrr::map(pluck(nest.gamm.M1, "gam"), ~summary(.))
[[1]]

Family: Negative Binomial(1.457)
Link function: log

Formula:
N ~ s(Site, bs = "re") + Generation + s(SessionDay, by = Generation,
   bs = "tp") + Company + s(Since.open, by = Company,
   bs = "tp")

Parametric coefficients:
             Estimate Std. Error z value Pr(>|z|)
(Intercept)  135.16506   29.83132   4.531 5.87e-06 ***
Generation1 -134.82759   29.84038  -4.518 6.23e-06 ***
Generation2 -134.53270   29.83091  -4.510 6.49e-06 ***
Generation3 -133.70707   29.83090  -4.482 7.39e-06 ***
CompanySC      0.76054    0.03772  20.165  < 2e-16 ***
CompanySE      1.51351    0.03685  41.076  < 2e-16 ***
---
Signif. codes:  0 ‘***’ 0.001 ‘**’ 0.01 ‘*’ 0.05 ‘.’ 0.1 ‘ ’ 1

Approximate significance of smooth terms:
                           edf Ref.df Chi.sq p-value
s(Site)                   2.979  3.000 426.73  <2e-16 ***
s(SessionDay):Generation0 8.097  8.237 745.32  <2e-16 ***
s(SessionDay):Generation1 6.238  6.610 772.70  <2e-16 ***
s(SessionDay):Generation2 8.475  8.921 708.74  <2e-16 ***
s(SessionDay):Generation3 8.687  8.971 797.49  <2e-16 ***
s(Since.open):CompanyOT   2.372  2.992  98.98  <2e-16 ***
s(Since.open):CompanySC   3.407  4.259  69.22  <2e-16 ***
s(Since.open):CompanySE   3.820  4.751  75.77  <2e-16 ***
---
Signif. codes:  0 ‘***’ 0.001 ‘**’ 0.01 ‘*’ 0.05 ‘.’ 0.1 ‘ ’ 1

R-sq.(adj) =   0.36   Deviance explained = 60.8%
-REML =  12615  Scale est. = 1         n = 4200

> # print AIC values
> nest.gamm.M2$aic
[1] 25072.31
> nest.gamm.M1$aic
[1] 25031.16
>
> nest.gamm.M2$aic-nest.gamm.M1$aic
[1] 41.14396


################################################################################################################################

# show the prediction for each model term

> unnest.gamm.M2[1, ] %>% select (intercept, starts_with("fit")) %>% data.frame() # also be starts_with("se")
  intercept      fit fit.Generation fit.Company fit.Since.open fit.Company.Since.open fit.s.Site.
1  135.7563 3.648757      -134.8113   0.5125618    -0.02564061             0.01109884   0.2173457
  fit.s.SessionDay..Generation0 fit.s.SessionDay..Generation1 fit.s.SessionDay..Generation2 fit.s.SessionDay..Generation3
1                             0                      1.988433                             0                             0
  fit.s.SessionDay..Generation
1                     1.988433

Interpretation of the M2 model

For each data row, the fitted value is obtained as the sum of:

fit ~   (intercept) +
        (fit.s.Site.) +
        (fit.Generation + fit.s.SessionDay..Generation) +
        (fit.Company + fit.Since.open + fit.Company.Since.open)


N                       Response variable, observation (1, 2, ...)
fit                     model prediction in the natural log scale.
resid.work              working residual used for the calculation of the partial residuals.

The predicted value for linear terms (the product of the parameter value and the coefficient)

intercept               (All companies, entire period) the size of intercept, taken from summary()
fit.Generation          (except for Generation 0) baseline density of male population in each season
fit.Company             (SC and SE, entire period) the difference in the lure attractiveness right after the exposure to the air.
                            Equal to CompanySC and CompanySE in summary().
fit.Since.open          (All companies, Since.open > 0) Daily change in the lure attractiveness.
fit.Company.Since.open  (SC and SE, Since.open > 0) Effect of the companies on the daily change in the lure attractiveness.

The predicted value for the (nonlinear) random effects

fit.s.Site.                     Difference in population density in each field in each generation. Centered to zero.
fit.s.SessionDay..Generation    Fluctuation in the male population density in each generation. Centered.
                                Assuming same shape in all fields in each generation.

############

> unnest.gamm.M1[1, ] %>% select (intercept, starts_with("fit")) %>% data.frame() # also be starts_with("se")
  intercept      fit fit.Generation fit.Company fit.s.Site. fit.s.SessionDay..Generation0 fit.s.SessionDay..Generation1
1  135.1651 3.335448      -134.8276   0.7605356    0.215987                             0                       2.01721
  fit.s.SessionDay..Generation2 fit.s.SessionDay..Generation3 fit.s.Since.open..CompanyOT fit.s.Since.open..CompanySC
1                             0                             0                           0                 0.004243302
  fit.s.Since.open..CompanySE fit.s.SessionDay..Generation fit.s.Since.open..Company
1                           0                      2.01721               0.004243302

(Note: fit.s.Since.open..Company is the sum of fit.s.Since.open..CompanyOT/SC/SE, calculated by hand)


Interpretation of the M1 model

fit ~   (intercept) +
        (fit.s.Site.) +
        (fit.Generation + fit.s.SessionDay..Generation) +
        (fit.Company + fit.s.Since.open..Company)

N                       Response variable, observation (1, 2, ...)
fit                     model prediction in the natural log scale.
resid.work              working residual used for the calculation of the partial residuals.

The predicted value for the lure attractiveness

intercept                   (All companies, entire period) the size of intercept, taken from summary()
fit.Generation          (except for Generation 0) baseline density of male population in each season
fit.Company                 (SC and SE, entire period) the difference in the lure attractiveness.
fit.s.Since.open..Company   Daily change in the lure attractiveness, having unique shape for every company and generation. Centered.

The predicted value for the (nonlinear) random effects

fit.s.Site.                     Difference in population density in each field in each generation. Centered to zero.
fit.s.SessionDay..Generation    Fluctuation in the male population density in each generation. Centered.
                                Assuming same shape in all fields in each generation.

################################################################################################################################


# Predict the fluctuation in the population size, without duplication of the data rows

# make the new data.frame for the high-resolution prediction of smoother terms: versus SessionDay, every 2hrs

# (if you want to avoid extrapolation of SessionDay for the generations 1 and 0, use the following)
dat.base.101 <- dat %>%
    dplyr::group_by(Generation) %>%
    tidyr::nest() %>%
    dplyr::mutate(  Minday=min(pluck(data, 1, "SessionDay")), Maxday=max(pluck(data, 1, "SessionDay"))  ) %>%
    dplyr::mutate(  SessionDay=list(seq(Minday, Maxday, by=1/12))  ) %>%
    dplyr::mutate(  data.101=purrr::map(.x=data, .f=. %>%
                                        dplyr::filter(  Site=="A", Company=="OT", Open==0, Since.open==1  ) %>%
                                        dplyr::select(  Year, Site, Company, Open, Open.str,
                                                        Since.open  ))  ) %>%
    dplyr::ungroup() %>%
    dplyr::select(-data) %>%
    tidyr::unnest(cols=c(data.101, SessionDay))
dat.base.101


# (This version is used for the final analysis, including extrapolation)
dat.base.101 <- dat %>%
    tidyr::nest(data=everything()) %>%
    dplyr::mutate(  Minday=min(pluck(data, 1, "SessionDay")), Maxday=max(pluck(data, 1, "SessionDay"))  ) %>%
    dplyr::mutate(  SessionDay=list(seq(Minday, Maxday, by=1/12))  ) %>%
    dplyr::mutate(  data.101=purrr::map(.x=data, .f=. %>%
                                        dplyr::filter(  Site=="A", Company=="OT", Open==0, Since.open==1  ) %>%
                                        dplyr::select(  Year, Generation, Site, Company, Open, Open.str,
                                                        Since.open  ) %>%
                                        tidyr::expand_grid(SessionDay=pluck(SessionDay, 1)))  ) %>%
    dplyr::ungroup() %>%
    dplyr::select(-data, -SessionDay) %>%
    tidyr::unnest(cols=c(data.101))
dat.base.101


# gratia::confint() is used to calculate the simultaneous confidence interval of the smoother
library(gratia)

# model M2: versus SessionDay (i.e., base density)
predict.M2.base.101 <- dat.base.101 %>%
    dplyr::mutate(Gen2=Generation) %>%
    dplyr::group_by(Gen2) %>%
    nest() %>%
    dplyr::bind_cols(select(ungroup(nest.gamm.M2), gam)) %>%
    dplyr::mutate(  coef=purrr::map(.x=gam, .f=~coef(.))  ) %>%
    dplyr::mutate(  intercept=purrr::map_dbl(.x=coef, .f=~pluck(., "(Intercept)"))  ) %>%
    dplyr::mutate(  pred.terms=purrr::map(  .x=gam,
                                            .f=~predict(., newdata=pluck(data, 1), type="terms", se.fit=TRUE) %>%
                                                data.frame() %>%
                                                as_tibble()  )  ) %>%
    dplyr::mutate(  simulconf.101=purrr::map(   .x=gam,
                                                .f=~confint(., parm="SessionDay", type="simultaneous",
                                                            newdata=data.frame(data)) %>%
                                                    dplyr::filter(Generation==Gen2) %>%
                                                    dplyr::rename(  confint.SessionDay=SessionDay,
                                                                    confint.Generation=Generation  )   )  ) %>%
    dplyr::select(-gam, -coef) %>%
    tidyr::unnest(cols=c(data, pred.terms, simulconf.101)) %>%
    dplyr::ungroup() %>%
    dplyr::mutate(  fit.s.SessionDay..Generation =  fit.s.SessionDay..Generation0+fit.s.SessionDay..Generation1+
                                                    fit.s.SessionDay..Generation2+fit.s.SessionDay..Generation3,
                    se.fit.s.SessionDay..Generation=se.fit.s.SessionDay..Generation0+se.fit.s.SessionDay..Generation1+
                                                    se.fit.s.SessionDay..Generation2+se.fit.s.SessionDay..Generation3  ) %>%
    dplyr::select(-Gen2, -Minday, -Maxday)
predict.M2.base.101
colnames(predict.M2.base.101)


# model M1: versus SessionDay (i.e., base density)

# Not used in the final version
predict.M1.base.101 <- dat.base.101 %>%
    dplyr::mutate(Gen2=Generation) %>%
    dplyr::group_by(Gen2) %>%
    nest() %>%
    dplyr::bind_cols(select(ungroup(nest.gamm.M1), gam)) %>%
    dplyr::mutate(  coef=purrr::map(.x=gam, .f=~coef(.))  ) %>%
    dplyr::mutate(  intercept=purrr::map_dbl(.x=coef, .f=~pluck(., "(Intercept)"))  ) %>%
    dplyr::mutate(  pred.terms=purrr::map(  .x=gam,
                                            .f=~predict(., newdata=pluck(data, 1), type="terms", se.fit=TRUE) %>%
                                                data.frame() %>%
                                                as_tibble()  )  ) %>%
    dplyr::mutate(  simulconf.101=purrr::map(   .x=gam,
                                                .f=~confint(., parm="SessionDay", type="simultaneous",
                                                            newdata=data.frame(data)) %>%
                                                    dplyr::filter(Generation==Gen2) %>%
                                                    dplyr::rename(  confint.SessionDay=SessionDay,
                                                                    confint.Generation=Generation  )   )  ) %>%
    dplyr::select(-gam, -coef) %>%
    tidyr::unnest(cols=c(data, pred.terms, simulconf.101)) %>%
    dplyr::ungroup() %>%
    dplyr::mutate(  fit.s.SessionDay..Generation =  fit.s.SessionDay..Generation0+fit.s.SessionDay..Generation1+
                                                    fit.s.SessionDay..Generation2+fit.s.SessionDay..Generation3,
                    se.fit.s.SessionDay..Generation=se.fit.s.SessionDay..Generation0+se.fit.s.SessionDay..Generation1+
                                                    se.fit.s.SessionDay..Generation2+se.fit.s.SessionDay..Generation3  ) %>%
    dplyr::select(-Gen2, -Minday, -Maxday)
predict.M1.base.101
colnames(predict.M1.base.101)


# make the new data.frame for the high-resolution prediction of smoother terms: versus Since.open, every 2hrs
dat.attractiveness.101 <- dat %>%
    dplyr::group_by(Company) %>%
    tidyr::nest() %>%
    dplyr::mutate(  Minday=min(pluck(data, 1, "Since.open")), Maxday=max(pluck(data, 1, "Since.open"))  ) %>%
    dplyr::mutate(  Since.open=list(seq(Minday, Maxday, by=1/12))  ) %>%
    dplyr::mutate(  data.101=purrr::map(.x=data, .f=. %>%
                                        dplyr::filter(  Generation==3, Site=="A", Open==0, SessionDay==1  ) %>%
                                        dplyr::select(  Year, Generation, Site, Open, Open.str,
                                                        SessionDay  ))  ) %>%
    dplyr::ungroup() %>%
    dplyr::select(-data) %>%
    tidyr::unnest(cols=c(data.101, Since.open))
dat.attractiveness.101

# versus Since.open (attenuation of lure attractiveness: only needed for nonlinear models)
predict.M1.attractiveness.101 <- dat.attractiveness.101 %>%
    dplyr::mutate(Gen2=Generation, Comp2=Company) %>%
    dplyr::group_by(Gen2, Comp2) %>%
    nest() %>%
    dplyr::bind_cols(select(ungroup(nest.gamm.M1), gam)) %>%
    dplyr::mutate(  coef=purrr::map(.x=gam, .f=~coef(.))  ) %>%
    dplyr::mutate(  intercept=purrr::map_dbl(.x=coef, .f=~pluck(., "(Intercept)"))  ) %>%
    dplyr::mutate(  pred.terms=purrr::map(  .x=gam,
                                            .f=~predict(., newdata=pluck(data, 1), type="terms", se.fit=TRUE) %>%
                                                data.frame() %>%
                                                as_tibble()  )  ) %>%
    dplyr::mutate(  simulconf.101=purrr::map(   .x=gam,
                                                .f=~confint(., parm="Since.open", type="simultaneous",
                                                            newdata=data.frame(data)) %>%
                                                    dplyr::filter(  Company==Comp2  ) %>%
                                                    dplyr::rename(  confint.Company=Company,
                                                                    confint.Since.open=Since.open  )   )  ) %>%
    dplyr::select(-gam, -coef) %>%
    tidyr::unnest(cols=c(data, pred.terms, simulconf.101)) %>%
    dplyr::ungroup() %>%
    dplyr::select(-Gen2, -Comp2, -Minday, -Maxday) %>%
    dplyr::mutate(  fit.s.Since.open..Company =     fit.s.Since.open..CompanyOT+fit.s.Since.open..CompanySC+
                                                    fit.s.Since.open..CompanySE,
                    se.fit.s.Since.open..Company =  se.fit.s.Since.open..CompanyOT+se.fit.s.Since.open..CompanySC+
                                                    se.fit.s.Since.open..CompanySE  )
predict.M1.attractiveness.101
colnames(predict.M1.attractiveness.101)


################################################################################################################################

# code for Figure 3


library(extrafont)
library(ggplot2)
library(ggthemes)
library(patchwork)
library(devEMF)

scale_company <- c( "OT"=hsv(343/360, 0.95, 0.82), "SC"=hsv(145/360, 0.95, 0.72), "SE"=hsv(208/360, 0.85, 0.65),
                    "LT"=hsv(0/360, 0, 0.1),
                    "Minus0w"=hsv(0.1, 0.3, 0.95), "Minus1w"=hsv(0.3, 0.3, 0.95),
                    "Minus2w"=hsv(0.5, 0.3, 0.95), "Minus3w"=hsv(0.7, 0.3, 0.95),
                    "Minus4w"=hsv(0.9, 0.3, 0.95) )
names_gen <- c( `1`="1st gen.",
                `2`="2nd gen.",
                `3`="3rd gen.",
                `0`="0th gen.",
                `2017`="2017",
                `2018`="2018" )

# M2, base population
YLIM <- log(c(1*10^-4, 1*10^4))
p17 <- ggplot(data=predict.M2.base.101)
p17 <- p17 +
    geom_hline(yintercept=log(seq(0.001, 0.01, by=0.001)), size=0.1, color=gray(0.85)) +
    geom_hline(yintercept=log(seq(0.01, 0.1, by=0.01)), size=0.1, color=gray(0.85)) +
    geom_hline(yintercept=log(seq(0.1, 1, by=0.1)), size=0.1, color=gray(0.85)) +
    geom_hline(yintercept=log(seq(1, 10, by=1)), size=0.1, color=gray(0.85)) +
    geom_hline(yintercept=log(seq(10, 100, by=10)), size=0.1, color=gray(0.85)) +
    geom_hline(yintercept=log(seq(100, 1000, by=100)), size=0.1, color=gray(0.85)) +
    geom_hline(yintercept=log(seq(1000, 10000, by=1000)), size=0.1, color=gray(0.85)) +
    geom_vline(xintercept=seq(0, 49, by=7), size=0.25, color=gray(0.85)) +
    geom_hline(yintercept=log(10^(seq(-3, 3, by=1))), size=0.25, color=gray(0.85)) +
    geom_point( data=unnest.gamm.M2,
                aes(y=resid.work*sqrt(weights.work) + intercept + fit.Generation + fit.s.SessionDay..Generation,
                    x=SessionDay,
                    group=interaction(Company, Generation, Site), color=Company, shape=Site), size=0.5, stroke=0.1 ) +
#    geom_ribbon(aes(ymin=lower, ymax=upper, x=SessionDay,
#                    group=interaction(Generation)   ), size=0.05, color="black", fill="white"  ) +
    geom_line(  aes(y=intercept + fit.Generation, x=SessionDay,
                    group=interaction(Generation) ), color="white", size=1  ) +
    geom_line(  aes(y=intercept + fit.Generation, x=SessionDay,
                    group=interaction(Generation) ), color="black", linetype="dashed", size=0.35  ) +
    geom_line(  aes(y=intercept + fit.Generation + fit.s.SessionDay..Generation, x=SessionDay,
                    group=interaction(Generation) ), color="white", size=1  ) +
    geom_line(  aes(y=intercept + fit.Generation + fit.s.SessionDay..Generation, x=SessionDay,
                    group=interaction(Generation) ), color="black", size=0.35  ) +
    facet_grid( ~ Year + Generation, margins=F, labeller=as_labeller(names_gen) ) +
    theme_classic(base_size=10) +
    scale_color_manual(name="Company", values=scale_company, guide=guide_legend(byrow=TRUE, nrow=1)) +
    scale_shape_manual(name="Site", values=1:4, guide=guide_legend(byrow=TRUE, nrow=1)) +
    theme(  panel.background=element_blank(),
            plot.background=element_rect(color=NA, fill=NA),
            plot.margin=unit(c(0, 0, 0.5, 0), "lines")  ) + # top, right, bottom, left
    theme(  text=element_text(family="Arial"),
            plot.title=element_blank(),
#            plot.title=element_text(family="Arial", face="bold", size=10, angle=0, hjust=0),
            strip.background=element_blank(),
            strip.text=element_text(family="Arial", face="bold", margin=margin(b=4, t=0, l=4, r=4, unit="pt")),
            axis.text.x=element_text(family="Arial Narrow", face="bold", size=8, angle=0, hjust=0.5),#
            axis.text.y=element_text(family="Arial Narrow", face="bold", size=8, angle=0, hjust=1)  ) + # hjust=0.5: center
    theme(  legend.position="top", legend.spacing=unit(0, "lines"),
            legend.margin=margin(b=0, t=0, l=4, r=5, unit="pt"), # the margin around each legend
            legend.box="horizontal", # align multiple legend series horizontally
            legend.box.margin=margin(b=0, t=1, l=1, r=2, unit="pt"), # margins around the full legend area
            legend.box.background=element_blank(), # do not show box around the legend
            legend.box.spacing=unit(4, unit="pt"), # space between legend box and the plot body
            legend.key.size=unit(15, "pt"), legend.key.height=unit(5, "pt"), # enlarge key but not increase margin
            legend.title=element_text(family="Arial", size=8), legend.text=element_text(family="Arial", face="bold")  ) +
    scale_x_continuous(expand=c(0, 1), limits=c(0, 21), breaks=seq(0, 21, by=7), labels=seq(0, 21, by=7) ) +
    scale_y_continuous(expand=c(0, 0), limits=YLIM, breaks=log(10^(seq(-3, 3, by=1))), labels=10^(seq(-3, 3, by=1))) +
    labs( title="A", x="Session day (trap settlement = 0)", y="Male density (arbitrary unit)" )
ggsave("Fig.M2.base.pdf", device=cairo_pdf, plot=p17, dpi=300, width=12/2.54, height=8/2.54)


# M2, fluctuation in capture attractiveness
YLIM <- log(c(0.1, 10))
p18 <- ggplot(data=unnest.gamm.M2 %>% filter(Site=="A"))
p18 <- p18 +
    geom_hline(yintercept=log(seq(0.1, 1, by=0.1)), size=0.2, color=gray(0.85)) +
    geom_hline(yintercept=log(seq(1, 10, by=1)), size=0.2, color=gray(0.85)) +
    geom_hline(yintercept=log(seq(10, 100, by=10)), size=0.2, color=gray(0.85)) +
    geom_vline(xintercept=seq(0, 49, by=7), size=0.2, color=gray(0.85)) +
#    geom_point( data=unnest.gamm.M2,
#                aes(y=resid.work*sqrt(weights.work) + fit.Company + fit.Since.open + fit.Company.Since.open,
#                    x=Since.open,
#                    group=interaction(Company, Generation, Site), color=Company, shape=Open.str), size=0.8, stroke=0.2 ) +
    geom_line(  aes(y=fit.Company + fit.Since.open + fit.Company.Since.open, x=Since.open,
                    color=Company, group=interaction(Company, Generation), linetype=Company), size=0.5  ) +
    coord_fixed(ratio=10) +
    theme_classic(base_size=10) +
    scale_fill_manual(name="Company", values=scale_company, guide=guide_legend(byrow=TRUE, nrow=2)) +
    scale_color_manual(name="Company", values=scale_company, guide=guide_legend(byrow=TRUE, nrow=2)) +
    scale_shape_manual(name="Company", values=1:5, guide=guide_legend(byrow=TRUE, nrow=2)) +
    theme(  panel.background=element_blank(),
            plot.background=element_rect(color=NA, fill=NA),
            plot.margin=unit(c(0, 0.5, 0, 0), "lines")  ) + # top, right, bottom, left
    theme(  text=element_text(family="Arial"),
            plot.title=element_blank(),
            strip.background=element_blank(),
            strip.text=element_text(family="Arial", face="bold", margin=margin(b=4, t=0, l=4, r=4, unit="pt")),
            axis.text.x=element_text(family="Arial Narrow", face="bold", size=8, angle=0, hjust=0.5),
            axis.text.y=element_text(family="Arial Narrow", face="bold", size=8, angle=0, hjust=1)  ) +
    theme(  legend.position="top", legend.spacing=unit(0, "lines"),
            legend.margin=margin(b=0, t=0, l=4, r=5, unit="pt"), # the margin around each legend
            legend.box="horizontal", # align multiple legend series horizontally
            legend.box.margin=margin(b=2, t=0, l=1, r=2, unit="pt"), # margins around the full legend area
            legend.box.background=element_blank(), # do not show box around the legend
            legend.box.spacing=unit(4, unit="pt"), # space between legend box and the plot body
            legend.key.size=unit(15, "pt"), legend.key.height=unit(5, "pt"), # enlarge key but not increase margin
            legend.title=element_text(family="Arial", size=8), legend.text=element_text(family="Arial", face="bold")  ) +
    scale_x_continuous(expand=c(0, 1), limit=c(0, 49), breaks=seq(0, 49, by=14), labels=seq(0, 49, by=14) ) +
    scale_y_continuous( expand=c(0, 0), limits=YLIM,
                        breaks=log(c(0.1, 1, 10, 100)), labels=c(0.1, 1, 10, 100) ) +
    labs( title="B", x="Lure exposure (days)", y="Lure attractiveness" )
ggsave("Fig.M2.attractiveness.pdf", device=cairo_pdf, plot=p18, dpi=300, width=8/2.54, height=8/2.54)


# M1, fluctuation in capture attractiveness
YLIM <- log(c(0.1, 10))
p19 <- ggplot(data=predict.M1.attractiveness.101)
p19 <- p19 +
    geom_hline(yintercept=log(seq(0.1, 1, by=0.1)), size=0.2, color=gray(0.85)) +
    geom_hline(yintercept=log(seq(1, 10, by=1)), size=0.2, color=gray(0.85)) +
    geom_hline(yintercept=log(seq(10, 100, by=10)), size=0.2, color=gray(0.85)) +
    geom_vline(xintercept=seq(0, 49, by=7), size=0.2, color=gray(0.85)) +
#    geom_point( data=unnest.gamm.M1,
#                aes(y=resid.work*sqrt(weights.work) + fit.Company + fit.s.Since.open..Company,
#                    x=Since.open,
#                    group=interaction(Company, Generation, Site), color=Company, shape=Open.str), size=0.8, stroke=0.2 ) +
    geom_line(  aes(y=fit.Company + fit.s.Since.open..Company,
                    x=Since.open, color=Company, group=interaction(Company, Generation), linetype=Company), size=0.5  ) +
    coord_fixed(ratio=10) +
    theme_classic(base_size=10) +
    scale_fill_manual(name="Company", values=scale_company, guide=guide_legend(byrow=TRUE, nrow=2)) +
    scale_color_manual(name="Company", values=scale_company, guide=guide_legend(byrow=TRUE, nrow=2)) +
    scale_shape_manual(name="Company", values=1:5, guide=guide_legend(byrow=TRUE, nrow=2)) +
    theme(  panel.background=element_blank(),
            plot.background=element_rect(color=NA, fill=NA),
            plot.margin=unit(c(0, 0.5, 0, 0), "lines")  ) + # top, right, bottom, left
    theme(  text=element_text(family="Arial"),
            plot.title=element_blank(),
            strip.background=element_blank(),
            strip.text=element_text(family="Arial", face="bold", margin=margin(b=4, t=0, l=4, r=4, unit="pt")),
            axis.text.x=element_text(family="Arial Narrow", face="bold", size=8, angle=0, hjust=0.5),
            axis.text.y=element_text(family="Arial Narrow", face="bold", size=8, angle=0, hjust=1)  ) +
    theme(  legend.position="top", legend.spacing=unit(0, "lines"),
            legend.margin=margin(b=0, t=0, l=4, r=5, unit="pt"), # the margin around each legend
            legend.box="horizontal", # align multiple legend series horizontally
            legend.box.margin=margin(b=2, t=0, l=1, r=2, unit="pt"), # margins around the full legend area
            legend.box.background=element_blank(), # do not show box around the legend
            legend.box.spacing=unit(4, unit="pt"), # space between legend box and the plot body
            legend.key.size=unit(15, "pt"), legend.key.height=unit(5, "pt"), # enlarge key but not increase margin
            legend.title=element_text(family="Arial", size=8), legend.text=element_text(family="Arial", face="bold")  ) +
    scale_x_continuous(expand=c(0, 1), limit=c(0, 49), breaks=seq(0, 49, by=14), labels=seq(0, 49, by=14) ) +
    scale_y_continuous( expand=c(0, 0), limits=YLIM,
                        breaks=log(c(0.1, 1, 10, 100)), labels=c(0.1, 1, 10, 100) ) +
    labs( title="C", x="Lure exposure (days)", y="Lure attractiveness" )
ggsave("Fig.M1.attractiveness.pdf", device=cairo_pdf, plot=p18, dpi=300, width=8/2.54, height=8/2.54)


pf13 <- p17 / (p18 | p19) +
#    plot_layout(widths=c(12, 6, 6)) +
#    plot_layout(heights=c(6, 6, 6)) +
    plot_annotation(tag_levels="a", tag_prefix="(", tag_suffix=")") &
    theme(plot.tag=element_text(size=12, face="bold"))
ggsave("Figure3.pdf", device=cairo_pdf, plot=pf13, dpi=300, width=12/2.54, height=12/2.54)
ggsave("Figure3.eps", device=cairo_pdf, plot=pf13, dpi=300, width=12/2.54, height=12/2.54)
ggsave("Figure3.png", type="cairo", plot=pf13, dpi=300, width=12/2.54, height=12/2.54)

emf(file="Figure3.emf", width=12/2.54, height=12/2.54)
print(pf13)
dev.off()


################################################################################################################################

################################################################################################################################


# Part III: Predict male density using the yield in the light trap of the corresponding day

## Ltlog: log(N.lt+0.5)


# model M18: linear for lure attractiveness, nonlinear for the coefficient of the light trap
ptime0 <- proc.time()
nest.gamm.M18 <- dat %>%
    tidyr::nest(data=everything()) %>%
    dplyr::mutate(  gam=purrr::map(.x=data, .f=~mgcv::gam(  N ~ s(Site, bs="re") +
                                                            Generation + s(Ltlog, by=Generation, bs="tp") +
                                                            Company*Since.open,
                                                            data=., family=nb(theta=NULL, link="log") ))  ) %>%
    dplyr::mutate(  aic=purrr::map_dbl(.x=gam, .f=~AIC(.))  ) %>%
    dplyr::mutate(  gcv=purrr::map_dbl(.x=gam, .f=~pluck(., "gcv.ubre"))  ) %>%
    dplyr::mutate(  coef=purrr::map(.x=gam, .f=~coef(.))  ) %>%
    dplyr::mutate(  intercept=purrr::map_dbl(.x=coef, .f=~pluck(., "(Intercept)"))  ) %>%
    dplyr::mutate(  deviance=purrr::map(  .x=gam, .f=~residuals(., type="deviance"))  ) %>%
    dplyr::mutate(  resid.work=purrr::map(  .x=gam, .f=~residuals(., type="working"))  ) %>%
    dplyr::mutate(  weights.work=purrr::map(.x=gam, .f=~pluck(., "weights"))  ) %>%
    dplyr::mutate(  pred.link=purrr::map(   .x=gam,
                                            .f=~as_tibble(data.frame(predict(., type="link", se.fit=TRUE)))   )  ) %>%
    dplyr::mutate(  pred.terms=purrr::map(  .x=gam,
                                            .f=~as_tibble(data.frame(predict(., type="terms", se.fit=TRUE)))  )  )
cal.time <- proc.time()-ptime0; print(cal.time);
nest.gamm.M18
purrr::map(pluck(nest.gamm.M18, "gam"), ~summary(.))

colnames(nest.gamm.M18)
unnest.gamm.M18 <- nest.gamm.M18 %>%
    dplyr::select(-gam, -aic, -gcv, -coef) %>%
    tidyr::unnest(c(data, intercept, deviance, resid.work, weights.work, pred.link, pred.terms)) %>%
    dplyr::ungroup() %>%
    dplyr::mutate(  fit.s.Ltlog..Generation =  fit.s.Ltlog..Generation0+fit.s.Ltlog..Generation1+
                                                    fit.s.Ltlog..Generation2+fit.s.Ltlog..Generation3,
                    se.fit.s.Ltlog..Generation=se.fit.s.Ltlog..Generation0+se.fit.s.Ltlog..Generation1+
                                                    se.fit.s.Ltlog..Generation2+se.fit.s.Ltlog..Generation3  )
unnest.gamm.M18
colnames(unnest.gamm.M18)


# model M17: nonlinear for both of the lure attractiveness and light trap
ptime0 <- proc.time()
nest.gamm.M17 <- dat %>%
    tidyr::nest(data=everything()) %>%
    dplyr::mutate(  gam=purrr::map(.x=data, .f=~mgcv::gam(  N ~ s(Site, bs="re") +
                                                            Generation + s(Ltlog, by=Generation, bs="tp") +
                                                            Company + s(Since.open, by=Company, bs="tp"),
                                                            data=., family=nb(theta=NULL, link="log") ))  ) %>%
    dplyr::mutate(  aic=purrr::map_dbl(.x=gam, .f=~AIC(.))  ) %>%
    dplyr::mutate(  gcv=purrr::map_dbl(.x=gam, .f=~pluck(., "gcv.ubre"))  ) %>%
    dplyr::mutate(  coef=purrr::map(.x=gam, .f=~coef(.))  ) %>%
    dplyr::mutate(  intercept=purrr::map_dbl(.x=coef, .f=~pluck(., "(Intercept)"))  ) %>%
    dplyr::mutate(  deviance=purrr::map(  .x=gam, .f=~residuals(., type="deviance"))  ) %>%
    dplyr::mutate(  resid.work=purrr::map(  .x=gam, .f=~residuals(., type="working"))  ) %>%
    dplyr::mutate(  weights.work=purrr::map(.x=gam, .f=~pluck(., "weights"))  ) %>%
    dplyr::mutate(  pred.link=purrr::map(   .x=gam,
                                            .f=~as_tibble(data.frame(predict(., type="link", se.fit=TRUE)))   )  ) %>%
    dplyr::mutate(  pred.terms=purrr::map(  .x=gam,
                                            .f=~as_tibble(data.frame(predict(., type="terms", se.fit=TRUE)))  )  )
cal.time <- proc.time()-ptime0; print(cal.time);
nest.gamm.M17
purrr::map(pluck(nest.gamm.M17, "gam"), ~summary(.))

colnames(nest.gamm.M17)
unnest.gamm.M17 <- nest.gamm.M17 %>%
    dplyr::select(-gam, -aic, -gcv, -coef) %>%
    tidyr::unnest(c(data, intercept, deviance, resid.work, weights.work, pred.link, pred.terms)) %>%
    dplyr::ungroup() %>%
    dplyr::mutate(  fit.s.Ltlog..Generation =  fit.s.Ltlog..Generation0+fit.s.Ltlog..Generation1+
                                                    fit.s.Ltlog..Generation2+fit.s.Ltlog..Generation3,
                    se.fit.s.Ltlog..Generation=se.fit.s.Ltlog..Generation0+se.fit.s.Ltlog..Generation1+
                                                    se.fit.s.Ltlog..Generation2+se.fit.s.Ltlog..Generation3  ) %>%
    dplyr::mutate(  fit.s.Since.open..Company =     fit.s.Since.open..CompanyOT+fit.s.Since.open..CompanySC+
                                                    fit.s.Since.open..CompanySE,
                    se.fit.s.Since.open..Company =  se.fit.s.Since.open..CompanyOT+se.fit.s.Since.open..CompanySC+
                                                    se.fit.s.Since.open..CompanySE,  )
unnest.gamm.M17
colnames(unnest.gamm.M17)


# model M22: linear for lure attractiveness, nonlinear for the coefficient of the light trap,
# lacking intercept of generation
ptime0 <- proc.time()
nest.gamm.M22 <- dat %>%
    tidyr::nest(data=everything()) %>%
    dplyr::mutate(  gam=purrr::map(.x=data, .f=~mgcv::gam(  N ~ s(Site, bs="re") +
                                                            s(Ltlog, by=Generation, bs="tp") +
                                                            Company + s(Since.open, by=Company, bs="tp"),
                                                            data=., family=nb(theta=NULL, link="log") ))  ) %>%
    dplyr::mutate(  aic=purrr::map_dbl(.x=gam, .f=~AIC(.))  ) %>%
    dplyr::mutate(  gcv=purrr::map_dbl(.x=gam, .f=~pluck(., "gcv.ubre"))  ) %>%
    dplyr::mutate(  coef=purrr::map(.x=gam, .f=~coef(.))  ) %>%
    dplyr::mutate(  intercept=purrr::map_dbl(.x=coef, .f=~pluck(., "(Intercept)"))  ) %>%
    dplyr::mutate(  deviance=purrr::map(  .x=gam, .f=~residuals(., type="deviance"))  ) %>%
    dplyr::mutate(  resid.work=purrr::map(  .x=gam, .f=~residuals(., type="working"))  ) %>%
    dplyr::mutate(  weights.work=purrr::map(.x=gam, .f=~pluck(., "weights"))  ) %>%
    dplyr::mutate(  pred.link=purrr::map(   .x=gam,
                                            .f=~as_tibble(data.frame(predict(., type="link", se.fit=TRUE)))   )  ) %>%
    dplyr::mutate(  pred.terms=purrr::map(  .x=gam,
                                            .f=~as_tibble(data.frame(predict(., type="terms", se.fit=TRUE)))  )  )
cal.time <- proc.time()-ptime0; print(cal.time);
nest.gamm.M22
purrr::map(pluck(nest.gamm.M22, "gam"), ~summary(.))


# print AIC values
nest.gamm.M18$aic
nest.gamm.M17$aic
nest.gamm.M22$aic

nest.gamm.M18$aic-nest.gamm.M17$aic


################################################################################################################################

> # print AIC values
> nest.gamm.M18$aic
[1] 26069.27
> nest.gamm.M17$aic
[1] 26043.87
> nest.gamm.M22$aic
[1] 26057.39

> nest.gamm.M18$aic-nest.gamm.M17$aic
[1] 25.39507


> nest.gamm.M18
# A tibble: 1 x 11
 data          gam       aic    gcv coef     intercept deviance   resid.work  weights.work pred.link      pred.terms
 <list>        <list>  <dbl>  <dbl> <list>       <dbl> <list>     <list>      <list>       <list>         <list>
1 <tibble [4,2~ <gam>  26069. 13122. <dbl [4~      3.60 <dbl [4,2~ <dbl [4,20~ <dbl [4,200~ <tibble [4,20~ <tibble [4,200~

> purrr::map(pluck(nest.gamm.M18, "gam"), ~summary(.))
[[1]]

Family: Negative Binomial(1.061)
Link function: log

Formula:
N ~ s(Site, bs = "re") + Generation + s(Ltlog, by = Generation,
   bs = "tp") + Company * Since.open

Parametric coefficients:
                      Estimate Std. Error z value Pr(>|z|)
(Intercept)            3.597605   0.475316   7.569 3.76e-14 ***
Generation1           -2.053785   1.117094  -1.839  0.06599 .
Generation2          -15.100048   3.436053  -4.395 1.11e-05 ***
Generation3            1.253137   8.768924   0.143  0.88636
CompanySC              0.485462   0.094455   5.140 2.75e-07 ***
CompanySE              1.224248   0.092941  13.172  < 2e-16 ***
Since.open            -0.032001   0.002772 -11.545  < 2e-16 ***
CompanySC:Since.open   0.011579   0.003722   3.110  0.00187 **
CompanySE:Since.open   0.009722   0.003655   2.660  0.00781 **
---
Signif. codes:  0 ‘***’ 0.001 ‘**’ 0.01 ‘*’ 0.05 ‘.’ 0.1 ‘ ’ 1

Approximate significance of smooth terms:
                      edf Ref.df Chi.sq p-value
s(Site)              2.971  3.000 305.77  <2e-16 ***
s(Ltlog):Generation0 3.247  3.704  82.19  <2e-16 ***
s(Ltlog):Generation1 5.934  6.158 440.39  <2e-16 ***
s(Ltlog):Generation2 7.740  7.954 511.34  <2e-16 ***
s(Ltlog):Generation3 6.959  7.024 599.84  <2e-16 ***
---
Signif. codes:  0 ‘***’ 0.001 ‘**’ 0.01 ‘*’ 0.05 ‘.’ 0.1 ‘ ’ 1

R-sq.(adj) =  0.332   Deviance explained = 49.2%
-REML =  13122  Scale est. = 1         n = 4200


> nest.gamm.M17
# A tibble: 1 x 11
  data          gam       aic    gcv coef     intercept deviance   resid.work  weights.work pred.link      pred.terms
  <list>        <list>  <dbl>  <dbl> <list>       <dbl> <list>     <list>      <list>       <list>         <list>
1 <tibble [4,2~ <gam>  26044. 13103. <dbl [7~      2.84 <dbl [4,2~ <dbl [4,20~ <dbl [4,200~ <tibble [4,20~ <tibble [4,200~

> purrr::map(pluck(nest.gamm.M17, "gam"), ~summary(.))
[[1]]

Family: Negative Binomial(1.07)
Link function: log

Formula:
N ~ s(Site, bs = "re") + Generation + s(Ltlog, by = Generation,
    bs = "tp") + Company + s(Since.open, by = Company,
    bs = "tp")

Parametric coefficients:
             Estimate Std. Error z value Pr(>|z|)
(Intercept)   2.83671    0.47818   5.932 2.99e-09 ***
Generation1  -2.08475    1.13349  -1.839   0.0659 .
Generation2 -15.09641    3.43968  -4.389 1.14e-05 ***
Generation3   1.27769    8.88679   0.144   0.8857
CompanySC     0.74356    0.04169  17.837  < 2e-16 ***
CompanySE     1.45244    0.04096  35.461  < 2e-16 ***
---
Signif. codes:  0 ‘***’ 0.001 ‘**’ 0.01 ‘*’ 0.05 ‘.’ 0.1 ‘ ’ 1

Approximate significance of smooth terms:
                          edf Ref.df Chi.sq p-value
s(Site)                 2.971  3.000 303.74  <2e-16 ***
s(Ltlog):Generation0    3.270  3.723  84.25  <2e-16 ***
s(Ltlog):Generation1    5.942  6.165 444.14  <2e-16 ***
s(Ltlog):Generation2    7.741  7.955 503.23  <2e-16 ***
s(Ltlog):Generation3    6.961  7.024 593.11  <2e-16 ***
s(Since.open):CompanyOT 1.006  1.011 134.54  <2e-16 ***
s(Since.open):CompanySC 3.513  4.387  92.72  <2e-16 ***
s(Since.open):CompanySE 2.226  2.810  84.67  <2e-16 ***
---
Signif. codes:  0 ‘***’ 0.001 ‘**’ 0.01 ‘*’ 0.05 ‘.’ 0.1 ‘ ’ 1

R-sq.(adj) =   0.34   Deviance explained = 49.6%
-REML =  13103  Scale est. = 1         n = 4200

> nest.gamm.M22
# A tibble: 1 x 11
  data          gam       aic    gcv coef     intercept deviance   resid.work  weights.work pred.link      pred.terms
  <list>        <list>  <dbl>  <dbl> <list>       <dbl> <list>     <list>      <list>       <list>         <list>
1 <tibble [4,2~ <gam>  26057. 13112. <dbl [7~     0.336 <dbl [4,2~ <dbl [4,20~ <dbl [4,200~ <tibble [4,20~ <tibble [4,200~


> purrr::map(pluck(nest.gamm.M22, "gam"), ~summary(.))
[[1]]

Family: Negative Binomial(1.067)
Link function: log

Formula:
N ~ s(Site, bs = "re") + s(Ltlog, by = Generation, bs = "tp") +
    Company + s(Since.open, by = Company, bs = "tp")

Parametric coefficients:
            Estimate Std. Error z value Pr(>|z|)
(Intercept)  0.33571    0.71002   0.473    0.636
CompanySC    0.74114    0.04172  17.765   <2e-16 ***
CompanySE    1.45446    0.04099  35.484   <2e-16 ***
---
Signif. codes:  0 ‘***’ 0.001 ‘**’ 0.01 ‘*’ 0.05 ‘.’ 0.1 ‘ ’ 1

Approximate significance of smooth terms:
                          edf Ref.df Chi.sq p-value
s(Site)                 2.971  3.000 307.60  <2e-16 ***
s(Ltlog):Generation0    5.374  5.650 102.95  <2e-16 ***
s(Ltlog):Generation1    6.363  6.608 452.60  <2e-16 ***
s(Ltlog):Generation2    7.008  7.362 466.34  <2e-16 ***
s(Ltlog):Generation3    7.940  8.011 649.92  <2e-16 ***
s(Since.open):CompanyOT 1.001  1.002 129.28  <2e-16 ***
s(Since.open):CompanySC 3.488  4.357  89.06  <2e-16 ***
s(Since.open):CompanySE 2.283  2.882  82.39  <2e-16 ***
---
Signif. codes:  0 ‘***’ 0.001 ‘**’ 0.01 ‘*’ 0.05 ‘.’ 0.1 ‘ ’ 1

R-sq.(adj) =   0.33   Deviance explained = 49.5%
-REML =  13112  Scale est. = 1         n = 4200

################################################################################################################################


# Predict the fluctuation in the population size, without duplication of the data rows

# make the new data.frame for the high-resolution prediction of smoother terms: versus Ltlog

# (if you want to avoid extrapolation of SessionDay for the generations 1 and 0, use the following)
dat.base.101.lt <- dat %>%
    dplyr::group_by(Generation) %>%
    tidyr::nest() %>%
    dplyr::mutate(  Minlt=min(pluck(data, 1, "Ltlog")), Maxlt=max(pluck(data, 1, "Ltlog"))  ) %>%
    dplyr::mutate(  Ltlog=list(seq(Minlt, Maxlt, length.out=201))  ) %>%
    dplyr::mutate(  data.101=purrr::map(.x=data, .f=. %>%
                                        dplyr::filter(  Site=="A", Company=="OT", Open==0, Since.open==1  ) %>%
                                        dplyr::select(  Year, Site, Company, Open, Open.str,
                                                        SessionDay, Since.open  ))  ) %>%
    dplyr::ungroup() %>%
    dplyr::select(-data) %>%
    tidyr::unnest(cols=c(data.101, Ltlog))
dat.base.101.lt

# (This version is used for the final analysis, including extrapolation)
dat.base.101.lt <- dat %>%
    tidyr::nest(data=everything()) %>%
    dplyr::mutate(  Minlt=min(pluck(data, 1, "Ltlog")), Maxlt=max(pluck(data, 1, "Ltlog"))  ) %>%
    dplyr::mutate(  Ltlog=list(seq(Minlt, Maxlt, length.out=201))  ) %>%
    dplyr::mutate(  data.101=purrr::map(.x=data, .f=. %>%
                                        dplyr::filter(  Site=="A", Company=="OT", Open==0, Since.open==1  ) %>%
                                        dplyr::select(  Year, Generation, Site, Company, Open, Open.str,
                                                        SessionDay, Since.open  ) %>%
                                        tidyr::expand_grid(Ltlog=pluck(Ltlog, 1)))  ) %>%
    dplyr::ungroup() %>%
    dplyr::select(-data, -Ltlog) %>%
    tidyr::unnest(cols=c(data.101))
dat.base.101.lt


# gratia::confint() is used to calculate the simultaneous confidence interval of the smoother
library(gratia)

# model M18: versus Ltlog (i.e., base density)
predict.M18.base.101 <- dat.base.101.lt %>%
    dplyr::mutate(Gen2=Generation) %>%
    dplyr::group_by(Gen2) %>%
    nest() %>%
    dplyr::bind_cols(select(ungroup(nest.gamm.M18), gam)) %>%
    dplyr::mutate(  coef=purrr::map(.x=gam, .f=~coef(.))  ) %>%
    dplyr::mutate(  intercept=purrr::map_dbl(.x=coef, .f=~pluck(., "(Intercept)"))  ) %>%
    dplyr::mutate(  pred.terms=purrr::map(  .x=gam,
                                            .f=~predict(., newdata=pluck(data, 1), type="terms", se.fit=TRUE) %>%
                                                data.frame() %>%
                                                as_tibble()  )  ) %>%
    dplyr::mutate(  simulconf.101=purrr::map(   .x=gam,
                                                .f=~confint(., parm="Ltlog", type="simultaneous",
                                                            newdata=data.frame(data)) %>%
                                                    dplyr::filter(Generation==Gen2) %>%
                                                    dplyr::rename(  confint.Ltlog=Ltlog,
                                                                    confint.Generation=Generation  )   )  ) %>%
    dplyr::select(-gam, -coef) %>%
    tidyr::unnest(cols=c(data, pred.terms, simulconf.101)) %>%
    dplyr::ungroup() %>%
    dplyr::mutate(  fit.s.Ltlog..Generation =  fit.s.Ltlog..Generation0+fit.s.Ltlog..Generation1+
                                                    fit.s.Ltlog..Generation2+fit.s.Ltlog..Generation3,
                    se.fit.s.Ltlog..Generation=se.fit.s.Ltlog..Generation0+se.fit.s.Ltlog..Generation1+
                                                    se.fit.s.Ltlog..Generation2+se.fit.s.Ltlog..Generation3  ) %>%
    dplyr::select(-Gen2, -Minlt, -Maxlt)
predict.M18.base.101
colnames(predict.M18.base.101)


# model M17: versus Ltlog (i.e., base density)
predict.M17.base.101 <- dat.base.101.lt %>%
    dplyr::mutate(Gen2=Generation) %>%
    dplyr::group_by(Gen2) %>%
    nest() %>%
    dplyr::bind_cols(select(ungroup(nest.gamm.M17), gam)) %>%
    dplyr::mutate(  coef=purrr::map(.x=gam, .f=~coef(.))  ) %>%
    dplyr::mutate(  intercept=purrr::map_dbl(.x=coef, .f=~pluck(., "(Intercept)"))  ) %>%
    dplyr::mutate(  pred.terms=purrr::map(  .x=gam,
                                            .f=~predict(., newdata=pluck(data, 1), type="terms", se.fit=TRUE) %>%
                                                data.frame() %>%
                                                as_tibble()  )  ) %>%
    dplyr::mutate(  simulconf.101=purrr::map(   .x=gam,
                                                .f=~confint(., parm="Ltlog", type="simultaneous",
                                                            newdata=data.frame(data)) %>%
                                                    dplyr::filter(Generation==Gen2) %>%
                                                    dplyr::rename(  confint.Ltlog=Ltlog,
                                                                    confint.Generation=Generation  )   )  ) %>%
    dplyr::select(-gam, -coef) %>%
    tidyr::unnest(cols=c(data, pred.terms, simulconf.101)) %>%
    dplyr::ungroup() %>%
    dplyr::mutate(  fit.s.Ltlog..Generation =  fit.s.Ltlog..Generation0+fit.s.Ltlog..Generation1+
                                                    fit.s.Ltlog..Generation2+fit.s.Ltlog..Generation3,
                    se.fit.s.Ltlog..Generation=se.fit.s.Ltlog..Generation0+se.fit.s.Ltlog..Generation1+
                                                    se.fit.s.Ltlog..Generation2+se.fit.s.Ltlog..Generation3  ) %>%
    dplyr::select(-Gen2, -Minlt, -Maxlt)
predict.M17.base.101
colnames(predict.M17.base.101)


# make the new data.frame for the high-resolution prediction of smoother terms: versus Since.open, every 2hrs
dat.attractiveness.101.lt <- dat %>%
    dplyr::group_by(Company) %>%
    tidyr::nest() %>%
    dplyr::mutate(  Minday=min(pluck(data, 1, "Since.open")), Maxday=max(pluck(data, 1, "Since.open"))  ) %>%
    dplyr::mutate(  Since.open=list(seq(Minday, Maxday, by=1/12))  ) %>%
    dplyr::mutate(  data.101=purrr::map(.x=data, .f=. %>%
                                        dplyr::filter(  Site=="A", Open==0, SessionDay==1, Generation==2  ) %>%
                                        dplyr::select(  Year, Generation, Site, Open, Open.str, Ltlog, SessionDay  ))  ) %>%
    dplyr::ungroup() %>%
    dplyr::select(-data) %>%
    tidyr::unnest(cols=c(data.101, Since.open))
dat.attractiveness.101.lt


# versus Since.open (nonlinear attenuation of lure attractiveness: M17 model only)
predict.M17.attractiveness.101 <- dat.attractiveness.101.lt %>%
    dplyr::mutate(Gen2=Generation, Comp2=Company) %>%
    dplyr::group_by(Gen2, Comp2) %>%
    nest() %>%
    dplyr::bind_cols(select(ungroup(nest.gamm.M17), gam)) %>%
    dplyr::mutate(  coef=purrr::map(.x=gam, .f=~coef(.))  ) %>%
    dplyr::mutate(  intercept=purrr::map_dbl(.x=coef, .f=~pluck(., "(Intercept)"))  ) %>%
    dplyr::mutate(  pred.terms=purrr::map(  .x=gam,
                                            .f=~predict(., newdata=pluck(data, 1), type="terms", se.fit=TRUE) %>%
                                                data.frame() %>%
                                                as_tibble()  )  ) %>%
    dplyr::mutate(  simulconf.101=purrr::map(   .x=gam,
                                                .f=~confint(., parm="Since.open", type="simultaneous",
                                                            newdata=data.frame(data)) %>%
                                                    dplyr::filter(  Company==Comp2  ) %>%
                                                    dplyr::rename(  confint.Company=Company,
                                                                    confint.Since.open=Since.open  )   )  ) %>%
    dplyr::select(-gam, -coef) %>%
    tidyr::unnest(cols=c(data, pred.terms, simulconf.101)) %>%
    dplyr::ungroup() %>%
    dplyr::select(-Gen2, -Comp2, -Minday, -Maxday) %>%
    dplyr::mutate(  fit.s.Since.open..Company =     fit.s.Since.open..CompanyOT+fit.s.Since.open..CompanySC+
                                                    fit.s.Since.open..CompanySE,
                    se.fit.s.Since.open..Company =  se.fit.s.Since.open..CompanyOT+se.fit.s.Since.open..CompanySC+
                                                    se.fit.s.Since.open..CompanySE  )
predict.M17.attractiveness.101
colnames(predict.M17.attractiveness.101)


################################################################################################################################


# Figure 4

scale_gen <- c( "1"=hsv(56/360, 1, 0.76), "2"=hsv(310/360, 0.91, 0.75), "3"=hsv(346/360, 0.75, 0.91), "0"=hsv(0.5, 1, 0.7) )


# M2  model, calibration with light trap data
YLIM <- log(c(1*10^-5, 1*10^5))
p27 <- ggplot(data=predict.M18.base.101)
p27 <- p27 +
    geom_hline(yintercept=log(seq(0.001, 0.01, by=0.001)), size=0.1, color=gray(0.85)) +
    geom_hline(yintercept=log(seq(0.01, 0.1, by=0.01)), size=0.1, color=gray(0.85)) +
    geom_hline(yintercept=log(seq(0.1, 1, by=0.1)), size=0.1, color=gray(0.85)) +
    geom_hline(yintercept=log(seq(1, 10, by=1)), size=0.1, color=gray(0.85)) +
    geom_hline(yintercept=log(seq(10, 100, by=10)), size=0.1, color=gray(0.85)) +
    geom_hline(yintercept=log(seq(100, 1000, by=100)), size=0.1, color=gray(0.85)) +
    geom_hline(yintercept=log(10^(seq(-3, 3, by=1))), size=0.25, color=gray(0.85)) +
    geom_point( data=unnest.gamm.M18,
                aes(y=resid.work*sqrt(weights.work) + intercept + fit.Generation + fit.s.Ltlog..Generation,
                    x=Ltlog, group=interaction(Company, Generation, Site), color=Generation, shape=Generation ),
                size=0.8, stroke=0.15 ) +
    geom_line(  aes(y=intercept + fit.Generation + fit.s.Ltlog..Generation,
                    x=Ltlog, group=interaction(Company, Generation, Site, Open)),
                color="white", size=1  ) +
    geom_line(  aes(y=intercept + fit.Generation + fit.s.Ltlog..Generation,
                    x=Ltlog, group=interaction(Company, Generation, Site, Open), color=Generation, linetype=Generation),
                size=0.4  ) +
    theme_classic(base_size=10) +
    scale_color_manual(name="Generation", values=scale_gen, guide=guide_legend(byrow=TRUE, nrow=2)) +
    scale_shape_manual(name="Generation", values=1:4, guide=guide_legend(byrow=TRUE, nrow=2)) +
    theme(  panel.background=element_blank(),
            plot.background=element_rect(color=NA, fill=NA),
            plot.margin=unit(c(0, 0.5, 0, 0), "lines")  ) + # top, right, bottom, left
    theme(  text=element_text(family="Arial"),
            plot.title=element_blank(),
#            plot.title=element_text(family="Arial", face="bold", size=10, angle=0, hjust=0),
            strip.background=element_blank(),
            strip.text=element_text(family="Arial", face="bold", margin=margin(b=2, t=0, l=4, r=4, unit="pt")),
            axis.text.x=element_text(family="Arial Narrow", face="bold", size=8, angle=0, hjust=0.5),
            axis.text.y=element_text(family="Arial Narrow", face="bold", size=8, angle=0, hjust=1)  ) + # hjust=0.5: center
    theme(  legend.position="top", legend.spacing=unit(0, "lines"),
            legend.margin=margin(b=0, t=0, l=4, r=5, unit="pt"), # the margin around each legend
            legend.box="horizontal", # align multiple legend series horizontally
            legend.box.margin=margin(b=2, t=1, l=1, r=2, unit="pt"), # margins around the full legend area
            legend.box.background=element_blank(), # do not show box around the legend
            legend.box.spacing=unit(4, unit="pt"), # space between legend box and the plot body
            legend.key.size=unit(15, "pt"), legend.key.height=unit(5, "pt"), # enlarge key but not increase margin
            legend.title=element_text(family="Arial"), legend.text=element_text(family="Arial", face="bold")  ) +
    scale_x_continuous( expand=c(0, 0), limit=log(c(0.4, 2500)),
                        breaks=log(10^(seq(-1, 4, by=1))), labels=10^(seq(-1, 4, by=1))) +
    scale_y_continuous(expand=c(0, 0), limits=YLIM, breaks=log(10^(seq(-3, 3, by=1))), labels=10^(seq(-3, 3, by=1))) +
    labs( title="A", x="Capture in light trap (N+0.5)", y="Predicted capture of \npheromone trap" )
ggsave("Fig.M18.base.pdf", device=cairo_pdf, plot=p27, dpi=300, width=8/2.54, height=8/2.54)


YLIM <- log(c(0.1, 10))
p28 <- ggplot(data=unnest.gamm.M18 %>% filter(Site=="A"))
p28 <- p28 +
    geom_hline(yintercept=log(seq(0.1, 1, by=0.1)), size=0.25, color=gray(0.85)) +
    geom_hline(yintercept=log(seq(1, 10, by=1)), size=0.25, color=gray(0.85)) +
    geom_hline(yintercept=log(seq(10, 100, by=10)), size=0.25, color=gray(0.85)) +
    geom_vline(xintercept=seq(0, 49, by=7), size=0.25, color=gray(0.85)) +
    geom_line(  aes(y=fit.Company + fit.Since.open + fit.Company.Since.open, x=Since.open,
                    color=Company, group=interaction(Company, Generation), linetype=Company), size=0.5  ) +
    coord_fixed(ratio=10) +
    theme_classic(base_size=10) +
    scale_fill_manual(values=scale_company, guide=guide_legend(byrow=TRUE, nrow=2)) +
    scale_color_manual(values=scale_company, guide=guide_legend(byrow=TRUE, nrow=2)) +
    scale_shape_manual(values=1:5, guide=guide_legend(byrow=TRUE, nrow=2)) +
    theme(  panel.background=element_blank(),
            plot.background=element_rect(color=NA, fill=NA),
            plot.margin=unit(c(0, 0.5, 0, 0), "lines")  ) + # top, right, bottom, left
    theme(  text=element_text(family="Arial"),
            plot.title=element_blank(),
            strip.background=element_blank(),
            strip.text=element_text(family="Arial", face="bold", margin=margin(b=4, t=0, l=4, r=4, unit="pt")),
            axis.text.x=element_text(family="Arial Narrow", face="bold", size=8, angle=0, hjust=0.5),
            axis.text.y=element_text(family="Arial Narrow", face="bold", size=8, angle=0, hjust=1)  ) +
    theme(  legend.position="top", legend.spacing=unit(0, "lines"),
            legend.margin=margin(b=0, t=0, l=4, r=5, unit="pt"), # the margin around each legend
            legend.box="horizontal", # align multiple legend series horizontally
            legend.box.margin=margin(b=2, t=0, l=1, r=2, unit="pt"), # margins around the full legend area
            legend.box.background=element_blank(), # do not show box around the legend
            legend.box.spacing=unit(4, unit="pt"), # space between legend box and the plot body
            legend.key.size=unit(15, "pt"), legend.key.height=unit(5, "pt"), # enlarge key but not increase margin
            legend.title=element_text(family="Arial", size=8), legend.text=element_text(family="Arial", face="bold")  ) +
    scale_x_continuous(expand=c(0, 1), limit=c(0, 49), breaks=seq(0, 49, by=14), labels=seq(0, 49, by=14) ) +
    scale_y_continuous( expand=c(0, 0), limits=YLIM,
                        breaks=log(c(0.1, 1, 10, 100)), labels=c(0.1, 1, 10, 100) ) +
    labs( title="B", x="Lure exposure (days)", y="Lure attractiveness" )
ggsave("Fig.M18.attractiveness.pdf", device=cairo_pdf, plot=p28, dpi=300, width=8/2.54, height=8/2.54)


YLIM <- log(c(0.1, 10))
p29 <- ggplot(data=predict.M17.attractiveness.101)
p29 <- p29 +
    geom_hline(yintercept=log(seq(0.1, 1, by=0.1)), size=0.25, color=gray(0.85)) +
    geom_hline(yintercept=log(seq(1, 10, by=1)), size=0.25, color=gray(0.85)) +
    geom_hline(yintercept=log(seq(10, 100, by=10)), size=0.25, color=gray(0.85)) +
    geom_vline(xintercept=seq(0, 49, by=7), size=0.25, color=gray(0.85)) +
    geom_line(  aes(y=fit.Company + fit.s.Since.open..Company,
                    x=Since.open, color=Company, group=interaction(Company, Generation), linetype=Company), size=0.5  ) +
    coord_fixed(ratio=10) +
    theme_classic(base_size=10) +
    scale_fill_manual(values=scale_company, guide=guide_legend(byrow=TRUE, nrow=2)) +
    scale_color_manual(values=scale_company, guide=guide_legend(byrow=TRUE, nrow=2)) +
    scale_shape_manual(values=1:5, guide=guide_legend(byrow=TRUE, nrow=2)) +
    theme(  panel.background=element_blank(),
            plot.background=element_rect(color=NA, fill=NA),
            plot.margin=unit(c(0, 0.5, 0, 0), "lines")  ) + # top, right, bottom, left
    theme(  text=element_text(family="Arial"),
            plot.title=element_blank(),
            strip.background=element_blank(),
            strip.text=element_text(family="Arial", face="bold", margin=margin(b=4, t=0, l=4, r=4, unit="pt")),
            axis.text.x=element_text(family="Arial Narrow", face="bold", size=8, angle=0, hjust=0.5),
            axis.text.y=element_text(family="Arial Narrow", face="bold", size=8, angle=0, hjust=1)  ) +
    theme(  legend.position="top", legend.spacing=unit(0, "lines"),
            legend.margin=margin(b=0, t=0, l=4, r=5, unit="pt"), # the margin around each legend
            legend.box="horizontal", # align multiple legend series horizontally
            legend.box.margin=margin(b=2, t=0, l=1, r=2, unit="pt"), # margins around the full legend area
            legend.box.background=element_blank(), # do not show box around the legend
            legend.box.spacing=unit(4, unit="pt"), # space between legend box and the plot body
            legend.key.size=unit(15, "pt"), legend.key.height=unit(5, "pt"), # enlarge key but not increase margin
            legend.title=element_text(family="Arial", size=8), legend.text=element_text(family="Arial", face="bold")  ) +
    scale_x_continuous(expand=c(0, 1), limit=c(0, 49), breaks=seq(0, 49, by=14), labels=seq(0, 49, by=14) ) +
    scale_y_continuous( expand=c(0, 0), limits=YLIM,
                        breaks=log(c(0.1, 1, 10, 100)), labels=c(0.1, 1, 10, 100) ) +
    labs( title="C", x="Lure exposure (days)", y="Lure attractiveness" )
ggsave("Fig.M17.attractiveness.pdf", device=cairo_pdf, plot=p29, dpi=300, width=8/2.54, height=8/2.54)


pf23 <- (p27 | p28 | p29) +
    plot_layout(widths=c(6, 6, 6)) +
    plot_annotation(tag_levels="a", tag_prefix="(", tag_suffix=")") &
    theme(plot.tag=element_text(size=12, face="bold"))
ggsave("Figure4.pdf", device=cairo_pdf, plot=pf23, dpi=300, width=18/2.54, height=7/2.54)
ggsave("Figure4.eps", device=cairo_pdf, plot=pf23, dpi=300, width=18/2.54, height=7/2.54)
ggsave("Figure4.png", type="cairo", plot=pf23, dpi=300, width=18/2.54, height=7/2.54)

emf(file="Figure4.emf", width=18/2.54, height=7/2.54)
print(pf23)
dev.off()


################################################################################################################################

################################################################################################################################
```
